# Genome evolution and between-host transmission of *Spiroplasma* endosymbiont in wild communities of Morpho butterflies

**DOI:** 10.1101/2024.02.22.581604

**Authors:** Jonathan Filée, Manuela López-Villavicencio, Vincent Debat, Gabin Rignault, Rachel Fourdin, Camilo Salazar, Karina Lucas Silva-Brandão, Patrick Blandin, André Victor Lucci Freitas, Carolina Pardo-Diaz, Violaine Llaurens

## Abstract

The evolution of endosymbiont genomes is likely influenced by the ecological interactions with their hosts. Here, we studied the evolution of *Spiroplasma* genomes, as well as their transmission patterns within and between *Morpho* butterflies sampled in the wild. *Spiroplasma* was detected in 4 out of 11 *Morpho* species studied and displayed a 3 times larger genome size as compared to *Spiroplasma* genomes documented in other hosts. This inflation in genome size is caused by massive and recent expansion of various mobile genetic elements and by the acquisition of new genes stemming from prophages. Interestingly, these new Spiroplasma genomes also revealed a peculiar evolution of toxin genes in plasmids that may enhance host resistance to parasites. Phylogenetic comparisons with *Spiroplasma* extracted from other plant and insect host suggest multiple independent colonization of Lepidoptera by *Spiroplasma*, and probable horizontal exchanges among distantly-related butterfly species occurring in South America. In contrast, resequencing data obtained for multiple populations of the two sister-species *M. helenor* and *M. achilles* living in sympatry over the majority of their distribution revealed an opposite prevalence (97% in *M. achilles* and 3% in *M. helenor*), suggesting low levels of transmission between these sympatric host-species. Reconciliation analysis of the phylogenetic relationships of mitochondrial genomes within *M. achilles* and *Spiroplasma* strains furthermore confirms predominant vertical transfers of the endosymbiont within species. Altogether, our results indicate persistent interactions between *Spiroplasma* symbiont and some *Morpho* species, as well as contrasted prevalence among sympatric host-species, consistent with an evolution of ecological interactions between the endosymbiont and its different hosts that may modify their genomic evolution.

## Introduction

Ecological relationships between bacteria and their host species generate selective pressures acting on their genomes (Wenergreen 2002). In turn, the evolution of genes in bacteria can modify their ecological interactions with their hosts (Ochman & Moran 2001). Intracellular bacteria, or endosymbionts, offer the opportunity to investigate feedbacks between genome evolution and ecological interactions. The endosymbiont lifestyle is often associated with profound changes in bacterial genomes (Wernegreen 2017): the prevalence of genetic drift in endosymbiont populations and the specialization to their host can induce genome erosion and progressive loss of metabolic functions. As the result, gene loss is frequently observed when comparing endosymbiont genomes to genomes of closely-related free-living species (Latorre & Manzano-Marín 2017). The mutation accumulation and genome decay through Muller’s ratchet is indeed documented as a specific feature of endosymbiotic bacteria (Moran 1996). Such reduction in the number of functional genes may in turn increase the extinction risk of endosymbiont populations (Bennett and Moran 2015), depending on the level of dependence and of specialization between endosymbionts and their hosts.

Long-term persistence of some endosymbionts has nevertheless been documented, raising the question of how ecological interactions with different hosts may limit genome decay (Naito and Pawlowska 2016). Microbial endosymbionts impact several life history traits of their hosts (Drew et al. 2019), affecting both their survival and reproductive success (Hurst 2017), with both positive and negative documented effects of host fitness. Endosymbionts may indeed induce positive effects as nutritional providers (Sudakaran et al. 2017) or as protective agents against pathogens (Ballinger and Perlman 2019; King 2019). On the other hand, different negative impacts of endosymbionts on the host fitness have been reported : in vertically-transmitted endosymbiont, sex-ratio distortion in the offspring of infected females is frequently observed, leading to so-called male-killing effect (Harumoto and Lemaitre 2018), therefore affecting host reproductive success and population dynamics. Because of the evolution of specialized resistance to different endosymbiont distorters, cytoplasmic incompatibilities - whereby host individuals carrying different endosymbiont strains might fail to reproduce - can emerge (Werren 1998)and further impair the reproductive success of their host.

The vertical transmission of endosymbionts from mother to offspring may lead to co-speciation and to congruent phylogenetic relationships between host and endosymbionts lineages (Downie & Gullan 2005). Nevertheless, horizontal transmission and host shift has been documented in a number of endosymbiont (Ahmed et al. 2023). Transmission of endosymbionts between species could result from occasional hybridization events occurring between closely-related species living in sympatry. Horizontal transfers between hosts may also happen through transmission via other species: for instance, some usually vertically-transmitted endosymbionts of aphids could be horizontally transmitted by parasitoid wasps (Kaech & Vorburger 2021). Trophic interactions may also contribute to such horizontal transmission: endosymbiont transmissions have indeed been documented among insect hosts feeding on the same plants (Gonella et al. 2015, Chrostek et al. 2017). As a result, transmission of endosymbionts is more likely to happen between sympatric host-species and might depend on the level of phylogenetic relatedness and ecological niche overlap in the host species (Russell et al. 2009). Studying the evolution of endosymbionts genomes in closely-related species living in sympatry may thus shed light on the interplay between genome evolution and ecological interactions with different hosts. Recent publications of large genomic datasets in insects now allow to better characterize the prevalence of these endosymbionts throughout arthropods (Medina et al. 2023) but, also, to investigate the evolution of these endosymbionts and the diversity of their ecological relationships with different hosts in the wild.

*Spiroplasma* are endosymbiotic bacteria observed in both plants and insects; they can be transmitted horizontally by ingestion or maternally inherited. They are associated with a large variety of hosts, and their genomes appear highly eroded with reduced metabolic capacities, high proportion of pseudogenes, and elevated evolutionary rates (Gerth et al. 2021; Liu et al. 2022). In *Drosophila*, the incidence of *Spiroplasma* among natural population is generally low (Watts et al. 2009; Haselkorn 2010), but in some cases, the fitness advantages brought to their hosts, such as protection against parasitic nematodes, is associated with a higher prevalence of this endosymbiont (Jaenike et al. 2010). The protection against pathogens brought by Spiroplasma probably stems from the presence of specific *toxin* genes in their genome. For instance, ribosome-inactivating protein toxins (RIPs) have been shown to enhance defense against parasitoid wasps and parasitic nematodes in some *Drosophila* species carrying *Spiroplasma* from the *citri* clade (Hamilton et al., 2016; Ballinger and Perlman, 2017). The evolution of such toxin genes may promote the persistence of vertically- transmitted *Spiroplasma* in insects, because of the positive effects of host phenotypes (Moore & Ballinger 2023). Investigating the evolution of toxin gene families in *Spiroplasma* carried by insects can thus shed light on the ecological interactions with their hosts.

Experimental infections show that vertically-transmitted *Spiroplasma* can be transmitted across species (Nakayama et al. 2015) but such transfer between host species can be constrained by the phylogenetic distance between different host species (Tinsley and Majerus 2007). However, the tempo and pattern of *Spiroplasma* transmission in natural communities remain to be investigated. Therefore, studying the evolution of *Spiroplasma* genomes in natural communities of insect hosts can now shed light on the feedbacks between bacterial genome evolution and ecological interactions with their hosts.

In Lepidoptera, the diversity and the impact of cytoplasmic endosymbionts on host phenotypes have been studied in a handfull of species: *Spiroplasma* and *Wolbachia* are the most frequently reported endosymbionts, with various effects on host fitness (Duplouy and Hornett 2018). Both have been shown to generate male killing and sex ratio distortion in different Lepidoptera species, such as *Acraea encedon*, *Hypolimnas bolina* or *Danaus chrysippus* (Nymphalidae) (Duplouy and Hornett 2018; Jiggins et al. 2000). However, the evolution of the Spiroplasma genomes carried by butterflies in the wild, as well as the transmission of these endosymbionts within and across sympatric species of butterflies remains largely uncharacterized. Here, we focus on the *Spiroplasma* of *Morpho* butterfly species to characterize their level of ecological specialization, as well as their transmission mode. The genus *Morpho* is composed of emblematic species from the Neotropical rainforests, where up to ten different species can be observed in sympatry in Amazonian lowlands and the Guiana shield (Blandin and Purser 2013). Studying endosymbiont genomes found in *Morpho* butterflies from the wild allows to test (1) how much endosymbionts are shared across closely *vs.* distantly related host species, (2) how their genomes, and more specifically their toxin genes, evolved in different hosts, and (3) how endosymbionts are transmitted within and among sympatric host species. We used whole-genome sequencing data from 11 *Morpho* species to study the evolution of endosymbiont genomes in closely- related hosts. We then investigated the prevalence of endosymbionts in different populations of two sister-species of *Morpho* living in sympatry to characterize the transmission of the endosymbionts within and between species.

## Materials and Methods

### Genus dataset

To identify the diversity of endosymbiont genomes occurring in butterflies from the genus *Morpho* species, we analyzed the sequencing data obtained using the PacBio HiFi methodology applied to specimens from 11 Morpho species: 8 Amazonian species (*M. marcus*, *M. eugenia*, *M. telemachus*, *M. hecuba*, *M. rhetenor*, *M. menelaus*, *M. deidamia*, *M. achilles*), one widespread species, occurring from the Amazon to the South Atlantic Forest (*M. helenor*) and 2 sympatric species ranging from western Ecuador to Central America (*M. amathonte*, *M. granadensis)*. A single individual was sequenced for each species. Note that within *M. telemachus* there are two sympatric morphs (with either blue or yellow wings), so we analyzed one individual per morph. This dataset including all sampled species is referred to as the *genus dataset*.

For each individual included in the *genus dataset*, the DNA extraction was carried out from the thorax muscles of a male individual using the Qiagen Genomic-tip 100/G kit, following supplier instructions. After DNA extraction, the sequencing library was prepared following the manufacturer’s instructions “Procedure and Checklist Preparing HiFi SMRTbell Libraries Using SMRTbell Express Template Prep Kit 2.0.” for *M. helenor*, *M. achilles* and *M. deidamia* and “Procedure and Checklist – Preparing whole genome and metagenome libraries using SMRTbell® prep kit 3.0” for the other species. Libraries were sequenced on several PacBio Sequel II SMRT cells with the adaptive loading method or by diffusion loading on a SequelII instrument (for additional details see (Bastide et al. 2023). The reads were assembled into contigs using Hifiasm (Cheng et al. 2021) using the option no purge (-l0) to avoid eventual over-purging symbiont sequences. We used also alternative assemblers as HiCanu (Nurk et al. 2020) and Flye (Kolmogorov et al. 2019). The mitochondrial genome for all *Morpho* species was assembled directly from the PacBio Hifi reads with Rebaler (https://github.com/rrwick/Rebaler). For the assembly of mitochondrial genomes of *M. helenor*, *M. achilles* and *M. deidamia* the mitochondrial genome of the closely related species *Pararge aegeria* (Nymphalidae: Satyrinae) was used as a reference (Bastide et al. 2023) while for the other eight species, we used the genome of *M. helenor* as a reference.

### Endosymbiont metagenomic assembly

To detect the presence of endosymbiont genomes in the *genus dataset*, we used Blobtools (Laetsch and Blaxter 2017) with Diamond as search engine (Buchfink et al. 2015) against the UniProt database using a local copy of the NCBI TaxID file for the taxonomic assignment of the first best hits. Minimap2 (Li 2018) was used for read mapping with the options -ax map-hifi. Endosymbiont contigs were extracted using seqtk (available at https://github.com/lh3/seqtk) and processed through the Dfast workflow (Tanizawa et al. 2018) to estimate statistics and taxonomic assignment. The completeness of the detected endosymbiont genomes was estimated with CheckM (Parks et al. 2015) with the corresponding gene sets. Endosymbiont genome annotations were carried out using PROKKA (Seemann 2014) with standard parameters, and the corresponding genetic codes. Whole genome alignments were created using the nucmer utility of the Mummer package (Marçais et al. 2018) with standard options. Structural variations were visualized using D-GENIES (Cabanettes and Klopp 2018).

### Gene content and gene phylogeny in Spiroplasma genomes

We aimed at distinguishing orthologous genes shared with previously published *Spiroplasma* genomes found in other hosts, from genes specific to *Spiroplasma* in *Morpho*. We downloaded a set of 62 *Spiroplasma* genome assemblies with comparable genome metrics (N50>100kb) from the NCBI Refseq Genomes FTP server 12/02/2022 version (ftp://ftp.ncbi.nlm.nih.gov/genomes/refseq). We used Ortho-Finder 2.5.4 (Emms and Kelly 2015) to infer orthologous genes in the 62 *Spiroplasma* genomes as well as the *M. achilles Spiroplasma* genome sAch identified in this study. Then, functional annotation of the orthologous was inferred using the BlastKOALA tool against the KEGG database (Kanehisa et al. 2016).

We then studied putative toxin genes either involved in host-protective phenotypes or host male killing in insects. We used the HMMER software (Mistry et al. 2013) seeded with protein sequences of each symbiont genome and the Pfam domain sequence alignments (Finn et al. 2014) corresponding to the known toxin genes in *Spiroplasma* involved in Male-killing phenotype [OTU (PF02338 and OTU-like cysteine protease)] and those involved in protective phenotypes [RIP (PF00161)] as databases. Domain architectures of all matching proteins were then computed using PfamScan (Mistry et al. 2007) and SIGNALP 6.0 (Teufel et al. 2022). Phylogenies of the toxin proteins were built by extracting and aligning the corresponding OTU domains. The phylogenetic trees were inferred using IQ-TREE v2.1.3 (Nguyen et al. 2015) by estimating the best substitution models using ModelFinder (Kalyaanamoorthy et al. 2017). Branch support was then assessed by performing 1000 replicates using UltraFast boostraps (Hoang et al. 2018).

### Phylogeny of Spiroplasma

A previously published set of 96 single-copy, non-recombinant orthologs from the *Spiroplasma* genomes (Gerth et al. 2021) was used to assess the phylogenetic relationships of these endosymbionts. Orthologs were identified using the best reciprocal BLASTP hits of each of the 96 protein sequences using the *Spiroplasma poulsonii* sMel gene sequences as seeds and lead to a dataset of 72 gene sequences present in all of the *Morpho Spiroplasma* genomes. Alignments were computed using MAFFT (Katoh et al. 2002) and manually corrected to exclude ambiguous regions and taxa with sequence similarity >99% were removed. Phylogenic analyses were carried out using IQ-TREE v2.1.3 (Nguyen et al. 2015), and genes were partitioned to estimate the best substitution models using ModelFinder (Kalyaanamoorthy et al. 2017). Branch supports were assessed by performing 1000 replicates using UltraFast boostraps (Hoang et al. 2018). The resulting trees were rooted using the *Spiroplasma* sequences belonging to the *ixodetis* clade in accordance with the literature.

### Prediction of mobile genetic elements in Spiroplasma genomes

Inserted sequences (ISs) were identified by querying the ISFinder database (Siguier et al. 2006) with protein sequences of each endosymbiont genome assemblies using BLAST with *e-value* ≤10e-10 (Altschul et al. 1990).

Plasmid sequences were identified using the Plasflow software (Krawczyk et al. 2018) and prophage regions were found using two methods: (1) a sequence-similarity search using PHASTER (Arndt et al. 2016), and (2) a *de novo* prediction using PhiSpy (Akhter et al. 2012). Predictions gathered from the two methods were then merged in a single file. Comparative genomics with other prokaryotic genomes were then computed using a set of 25,674 genome sequences with comparable genome metrics (N50>100kb) downloaded from the NCBI Refseq Genomes FTP server 12/02/2022 version (ftp://ftp.ncbi.nlm.nih.gov/genomes/refseq). The corresponding proteomes were then downloaded and ISs were identified using the same procedure as described above for *Morpho* endosymbionts (using the 62 *Spiroplasma* genomes for which plasmids and phage regions were identified with the same previous workflow). Finally, CRISPR loci were identified using CRISPRFinder (Grissa et al. 2007).

### Sister-species dataset

To study the vertical and horizontal transmission of *Spiroplasma*, we focused on multiple populations of *M. achilles* and *M. helenor*, two sister-species living in sympatry across most of their distribution (Blandin and Purser 2013). We analyzed re-sequencing data obtained from 43 males of *M. helenor* and 33 individuals (30 males and 3 females) of *M. achilles* (Supplementary Table 1). The bias in the sex-ratio of the dataset is a direct consequence of the difficulties to collect *Morpho* females in the wild compared to males. This second dataset is referred to as the *sister-species dataset*.

### DNA extractions and genome sequencing of M. helenor and M. achilles

For the *sister-species dataset*, DNA for each individual was extracted from thorax muscle using the DNeasy Blood & Tissue Kit following the producer instructions. In most cases, DNA was extracted from SNAP-frozen individuals or samples preserved in DMSO, but we also used 13 samples of dried pinned *M. achilles* from the personal collection of Patrick Blandin (Supplementary Table 1).

Sequencing was then performed at the GeT-PlaGe core facility of INRAE. DNA-seq libraries were prepared using the Illumina TruSeq Nano DNA LT Library Prep Kit, following supplier instructions. Briefly, DNA was fragmented by sonication and adaptors were ligated. Eight cycles of PCR were then applied to amplify libraries. Library quality was assessed using an Advanced Analytical Fragment Analyzer and quantified by QPCR using the Kapa Library Quantification Kit. Sequencing was performed on an Illumina Novaseq 6000, using a paired-end read length of 2x150 pb on a S4 Flowcell.

### Population genomics of Spiroplasma in the sister-species dataset

Adaptors were removed from the reads with cutadapt (Martin 2011) and the reads of each individual were mapped against the original *Spiroplasma* genome found in the reference genome of *M. achilles* (referred to as *sAch* hereafter) using BWA-mem v0.7.17 (Li 2013). Then we used Samtools v1.10 (Li et al. 2009) to sort the resulting SAM and BAM files and recover the fasta sequences from *Spiroplasma* endosymbionts, which were then assembled using Megahit (Li et al. 2015) with default parameters. To assess the presence/absence of *Spiroplasma* in each individual, we blasted each contig against the *sAch* genome using BLASTN with e-value cuttoff = 10e-50 and identity percentage >90%. Matching contigs were extracted and the presence/absence of the endosymbiont in a sample was classified as follows: (i) ‘presence’ when more than 100kb of aligned matching contigs were obtained, (ii) ‘ambiguous’ when 5-100kb of aligned contigs were obtained, (iii) ‘absence’ when less than 5kb of aligned sequences were obtained. To validate this classification, we identified the 16S rDNA gene from each sample using BLASTN against the 16S rDNA gene from the reference genome *sAch*, with e-value cutoff of 10e-50 and identity percentage >90%. Then, we estimated a phylogenetic tree for *Spiroplasma* using the 16S rDNA sequences retrieved from the different butterfly samples, following the same procedure stated earlier.

Finally, RIP/Spaid toxin genes were also retrieved from each butterfly sample by applying a TBLASTN using the RIP/Spaid toxin gene identified in the sAch genome as seed, with e- value cuttoff = 10e-10.

### Tree reconciliation analysis used to test for host lateral switches

*Spiroplasma,* as maternally transmitted endosymbionts, are inherited together with the mitochondrial genome of the host. Therefore, we used ecceTERA (Jacox et al. 2016) to reconcile the mtDNA phylogeny of *Morpho* with the *Spiroplasma* phylogeny. The endosymbiont tree was obtained using a set of 133 marker genes computed with CheckM (Parks et al. 2015) Identical sequences were then removed to eliminate tree polytomies. SylvX (Chevenet et al. 2016) was used to visualize and interpret the reconciliation tree.The *Morpho* phylogeny was estimated with whole mitochondrial genomes, for which we combined those obtained in the genus-dataset and the sister-species datasets. Mitochondrial genomes for all individuals in the sister-species dataset were extracted directly from Illumina reads with GetOrganelle v1.7.5.3 (Jin et al. 2020) and the parameters -R 10 -k 21,45,65,85,105 -F animal_mt. Alignments were generated using MAFFT (Katoh et al. 2002) and manually curated to exclude ambiguous regions. The phylogeny was obtained with IQ-TREE v2.1.3 (Nguyen et al. 2015), estimating the best substitution model with ModelFinder (Kalyaanamoorthy et al. 2017), and assessing branch support with 1000 UltraFast boostrap replicates (Hoang et al. 2018). We used the *Heliconius melpomene* (Nymphalidae: Heliconiinae) sequence (NCBI accession HE579083) as an outgroup for the host tree and *Spiroplasma* strain sNigra for the symbiont tree (NCBI accession GCA_003987485.1).

## Results

### Morpho butterflies are sporadically associated with Spiroplasma symbionts

We sequenced and surveyed genomes of 11 species of *Morpho* butterflies (including two different morphs of *M. telemachus*), for the presence of endosymbiotic bacteria (so-called *genus dataset*). Contigs were binned using their GC%, read coverage and taxonomic assignation (Figure 1 and Supplementary Figure 1) ). The resulting blob-plots indicate the presence of a limited number of symbionts: the genomes of *M. achilles*, *M. amathonte* and *M. rhetenor* had contigs with very low GC% that match with *Spiroplasma*, whereas the genomes of *M. hecuba* and *M. helenor* had contigs associated with *Wolbachia* and *Enteroccocus*. In contrast, the genomes of the remaining six species of *Morpho* (i.e. *M. marcus*, *M. eugenia*, *M. telemachus*, *M. menelaus, M. deidamia* and *M. granadensis)* do not seem to harbor any symbiont sequences (Supplementary Figure 1).

**Figure 1:**
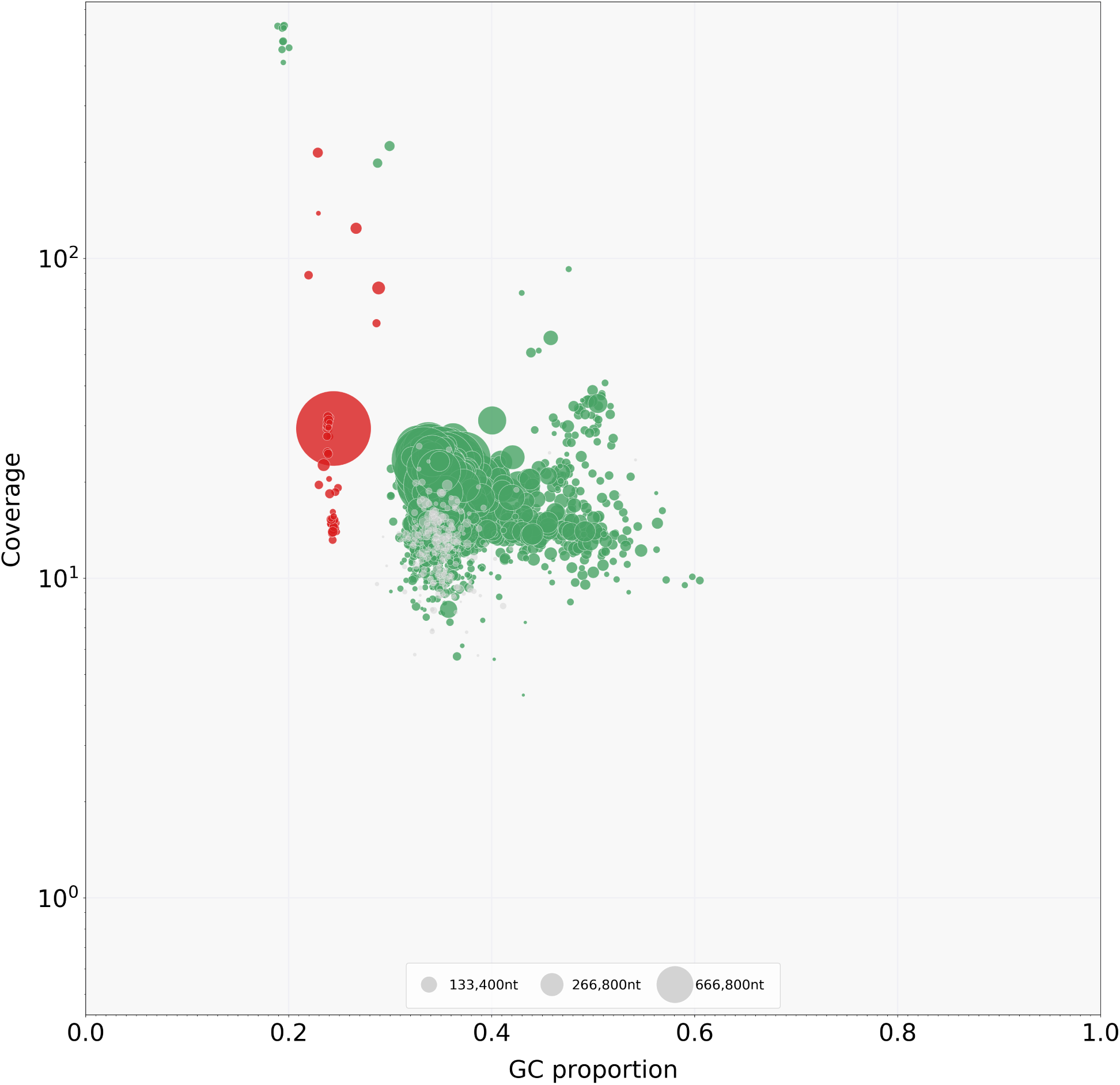
**Detection of symbionts within the genomic sequences of *Morpho achilles.*** Contigs represented as circles were binned based on their GC%, read coverage, and taxonomic assignment. Dark green contigs matched arthropod sequences and red contigs matched Mollicutes (*Spiroplasma*).The size of the circle is proportional to the size of the contigs.

We recovered and assembled three *Spiroplasma* genomes (*sAma*, *sAch* and *sRhe)* from the genomes of *M. amathonte*, *M. achilles* and *M. rhetenor*, respectively. These genomes are characterized by a very low GC% compared to their corresponding host genomes (24% versus 38%) and significant different read coverages (Figure 1). The *sAch* assembly was the most complete, containing 98% of a set of lineage-specific, single-copy, *Spiroplasma* marker genes (Table 1). The other two (*sAma* and *sRhe*) had lower completeness (60% and 78%,respectively, Table 1), indicating that only a fraction of the corresponding genomes was captured.

**Table 1.**
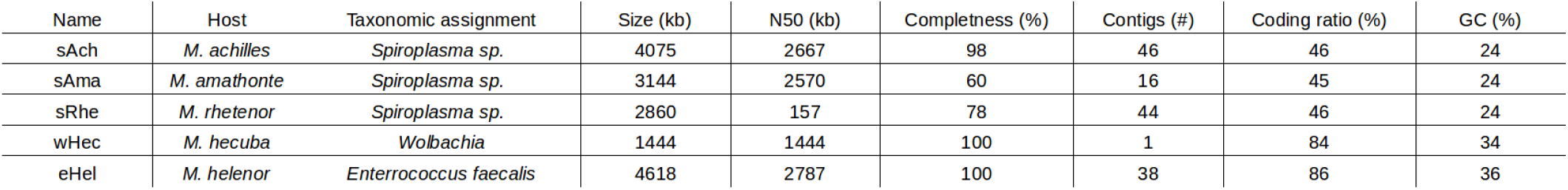
Assembly statistics of the endosymbiont genomes identified in genomes of *Morpho* butterflies.

### Inflation of genome size of Spiroplasma found in Morpho butterflies

The assemblies of *Spiroplasma* obtained from long-read data performed on *Morpho* butterfly samples display a considerably larger genome size (2,9Mb to 4,1Mb) than the 62 previously published genomes of this endosymbiont obtained with similar sequencing technics (1.1 Mb to 1.9 Mb; Supplementary Figure 2). We used alternative assembly softwares that also generate large *Spiroplasma* assemblies (up to 4.1Mb)(Supplementary Table 2). The sAch and sAma assemblies contained a large contig of 2,7 Mb and 2,5 Mb, respectively, with low levels of synteny (Supplementary Figure 3). The sAch and sAma assemblies were also composed of 46 and 15 small contigs ranging from 17kb to 74kb respectively, while the assembly of sRhe is composed of 44 contigs ranging from 14kb to 320kb (Figure 1). The 46 small contigs of the sAch assembly fall into 4 clusters based on sequence alignments, but all of them differ in size and/or in nucleotide similarity (Supplementary Figure 3).

To assess how much new genes contributed to the expansion of genome size in the *Spiroplasma* associated with *Morpho,* we searched for orthologous genes in sAch, which is the most complete assembly. Although the sAch genome encodes much more genes than others *Spiroplasma* genomes (5000 vs 1000-2000 genes) it contains a comparable number of conserved orthologous group of genes (ranging from 600 to 1000 ortho-groups). It has also encode for an unusually large number of species-specific genes absent in the other *Spiroplasma* genomes (>350 singletons; Supplementary Figure 4).

### Phylogeny of the Spiroplasma found in Morpho butterflies

To investigate the evolutionary origin of the *Spiroplasma* detected in *Morpho*, we built a phylogeny of this endosymbiont using a set of 72 concatenated single-copy genes present in all *Spiroplasma* genome assemblies with similar quality as the genomes obtained in this study, with recognizable homologs in the sAch complete genome. The assemblies *sAma*, *sAch* and *sRhe* retrieved from *Morpho* were monophyletic and are included within the *citri* clade (Figure 2), which includes diverse plant pathogens and endosymbionts of insects such as Hemiptera (*Spirolasma kunkelli*), Diptera (*S. sp. s*Nigra) and Hymenoptera (*S. melliferum*).

**Figure 2:**
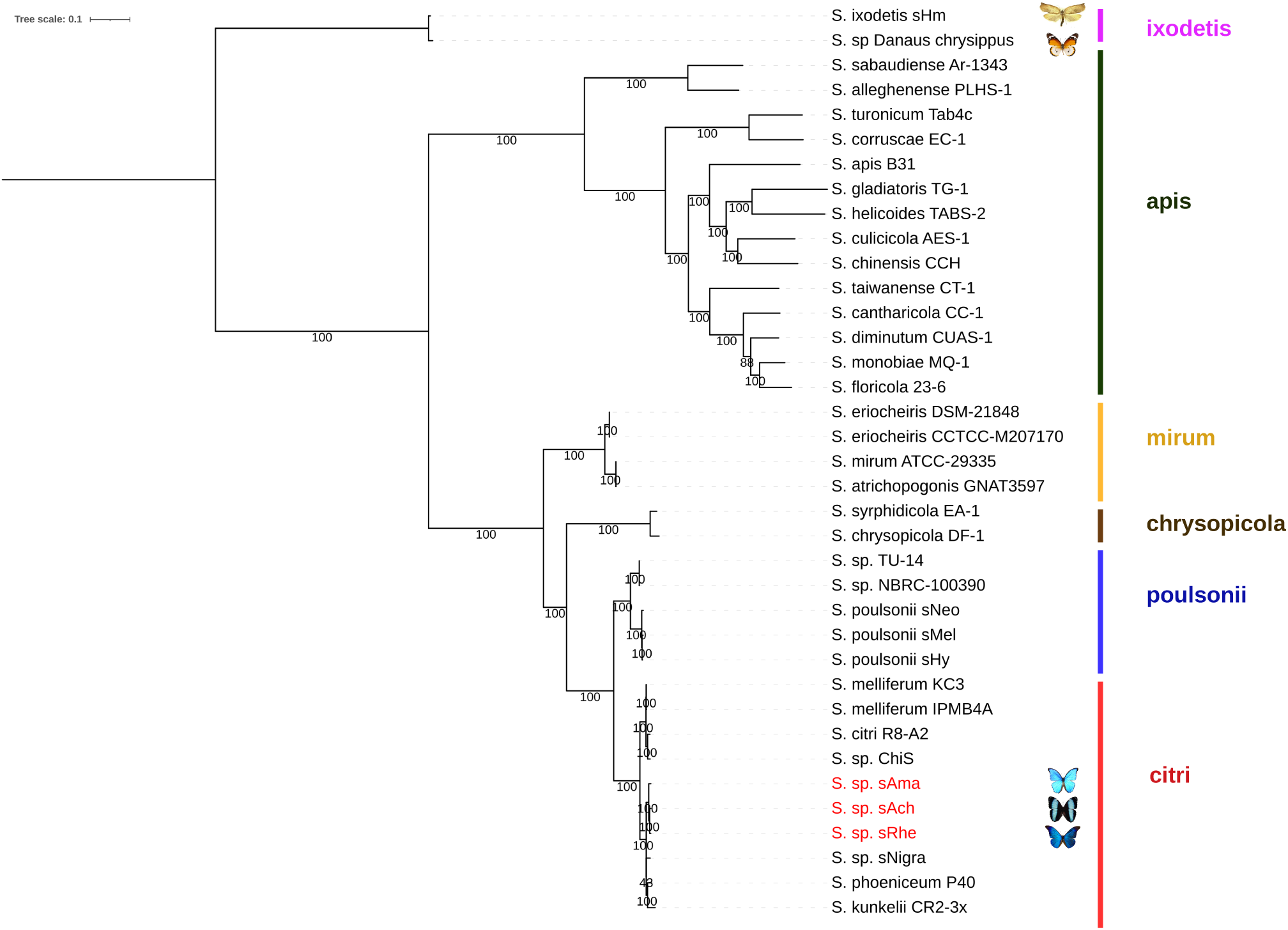
Phylogenetic reconstruction of *Spiroplasma* from different hosts based on the conserved set of 72 single copy orthologous genes. Recognized *Spiroplasma* clades are indicated in colors (right), *Spiroplasma* infecting butterflies are highlighted with pictures and the *Spiroplasma* detected in *M. amathonte*, *M. achilles* and *M. rhetenor* (sAma, SAch and sRhe, respectively) are highlighted in red. Phylogeny was constructed using 1000 bootstrap replicates. The phylogeny made with an extended gene dataset of 96 orthologs including only sAch produces the same topology (Supplementary Figure 5). Note that the *Spiroplasma* associated with the other Lepidoptera (the butterfly *Danaus chrysippus* and the moth *Homona magnanima*) fall in the distantly- related ixodetis clade (pink).

The 16S rDNA phylogeny that includes a broader taxonomic dataset confirms this observation (Supplementary Figure 6). The *Spiroplasma* recovered from *Morpho* are highly divergent from the *Spiroplasma* genomes found in other Lepidoptera such as the moth *Homona magnanima* (Tortricidae) (*S. ixodetis* sHM) or the nymphalid butterfly *Danaus chrysippus* (Nymphalidae: Danainae) (*S. sp. Danaus chrysipus*), both in the *ixodetis* clade (Figure 2). However, a *Spiroplasma* retrieved from the genome shotgun sequence of a South-American Hesperiid butterfly were found to belong to the *citri* clade as well (Moore & Ballinger 2023). This Hesperid symbiont found in *Jemadia suekentonmiller (*sSue) fall as sister taxa of the *Morpho Spiroplasma* in our phylogeny (Supplementary Figure 7). Taken together, these results suggest multiple independent colonization of Lepidoptera by *Spiroplasma*, and potential exchanges across South American butterfly species.

### Proliferation of mobile genetic elements in Spiroplasma found in Morpho

We found multiple mobile genetic elements (i.e., prokaryotic transposons, insertion sequences (IS), plasmids and prophages) integrated in the large genomes of the *Spiroplasma* retrieved from *Morpho.* We observed that IS are unusually abundant in these endosymbiont genomes, reaching a record-level in prokaryotes, which ranges from 398 to 885 copies (Figure 3A). The sAch assembly suggests this proliferation is associated with a surprisingly low number of IS families (Figure 3B). Indeed, only four IS families have expanded: an IS3-like family with 68 complete copies, an IS30 family with 96 complete copies, and an IS481 family with 72 intact copies. The fourth group is a 22kb composite transposon that we named Tn_sAch. This transposon has two IS3 copies at the tips and 24 conserved ORFs in the middle. In the assembly sAch, we observed Tn_sAch elements being especially common in the small contigs (constituting ∼54% of their length) and much less frequent in the large 2,7Mb contig (accounting for only 9% of its length). The strong structural conservation of the backbone of the 8 intact copies of Tn_sAch suggests *en bloc* successive transpositions in the genome.

**Figure 3:**
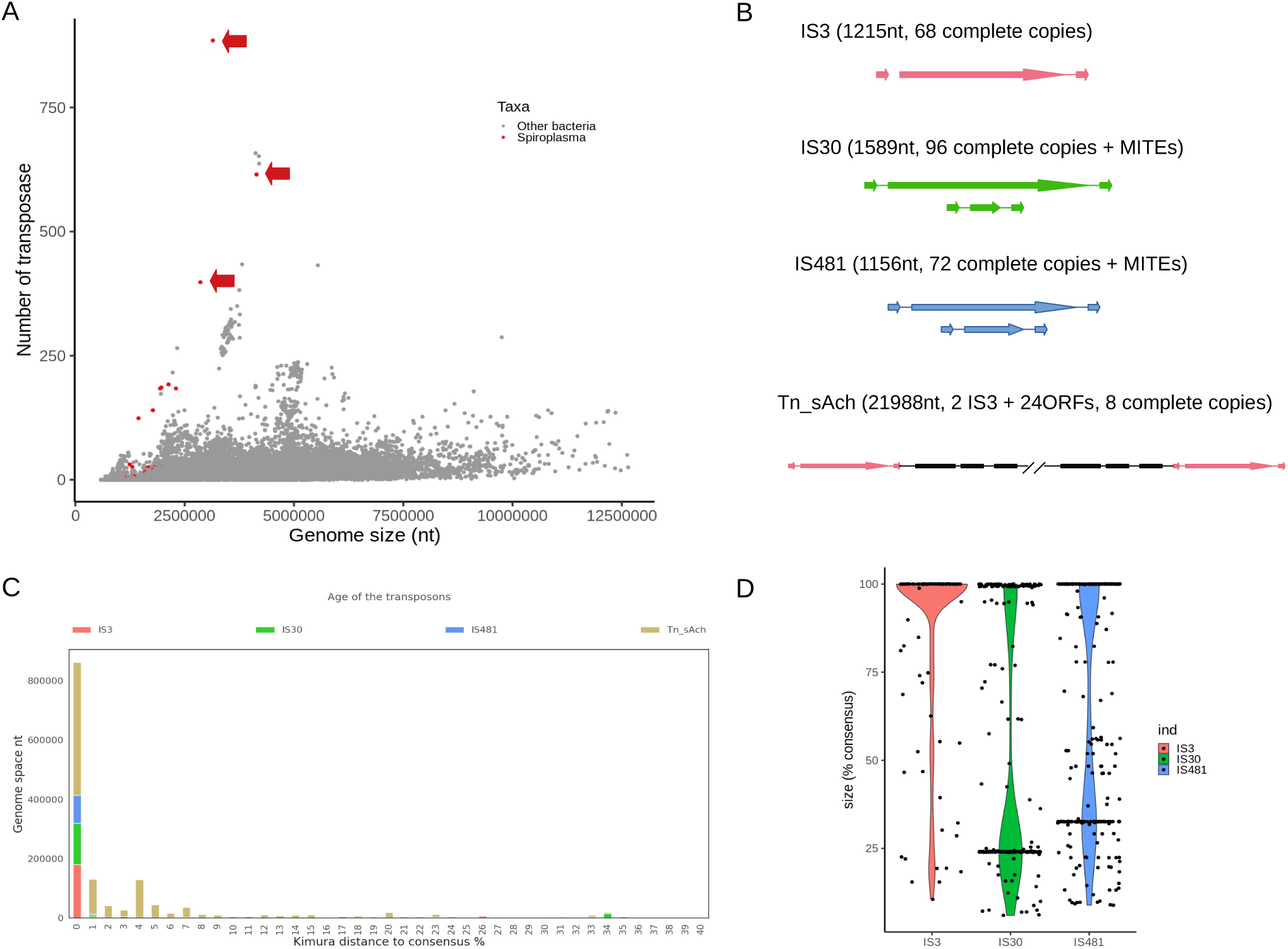
Insertion Sequences (ISs) found in the genomes of *Spiroplasma* found in *Morpho*. **A**: Number of transposase encoding genes found in a set of 25,675 prokaryotic genomes that include 62 *Spiroplasma* genomes (red dots) and three *Morpho Spiroplasma* (red dots with arrows) plotted against their genome sizes. **B**: Structure of the IS families found in the complete sAch genome and their main properties. Each colors represent a distinct IS families, arrows correspond to transposase genes and their internally deletted derivatives (MITEs), black rectangles indicate passenger genes of the compostite transposon. **C**: Analysis of the age of the IS copies in the sAch genome using the Kimura 2-parameter distance between the consensus sequence of a given family and all the individual copies that compose the family. The results are ordered based on the total amount of nucleotides. **D**: Analysis of the completeness of the different IS copies found in the sAch genome estimated as the percentage of the total length of the corresponding consensus sequences.

Sequence similarities of the different transposon copies analyzed as a proxy of the age of the different transposition events, indicates that most of them are identical or nearly identical, suggesting very recent transpositions (Figure 3C), but the copies of Tn_sAch have higher sequence divergence suggesting the presence of older copies. The IS30 and IS481 transposons have generated non-autonomous Miniature Inverted repeat Transposable Elements (MITEs) by internal deletion leading to smaller transposons that represent 25% and 32% of the size of the parental elements (Figure 3D). In contrast, most of the complete autonomous IS copies are 100% full-length and presumably intact, showing few truncated copies. Such high level of complete and identical IS copies strongly suggests that these families have recently expanded in the genomes of the *Spiroplasma* found in *Morpho* butterflies.

Our analyses of phage sequence invasion reveal the 28 to 32 integrated prophages in the genomes of *Spiroplasma* in *Morpho,* accounting for 418 kb in sAma, 482 kb in sRhe, and 538 kb in sAch (Supplementary Figure 8). The density of phage-derived elements (7.8/Mb) is thus larger than that in the most prophage-rich bacterial genome known to date (6.9/Mb) (Touchon et al. 2016; Frost et al. 2020). Interestingly, more than 80% of the sAch species-specific genes (singletons) are located in phage regions (Supplementary Figure 4). Functional analysis of these genes indicated that they are enriched by two KEGG functional categories: “genetic information processing” and “signaling and cellular processes” (Supplementary Figure 4). Therefore, the genome size expansion in *Spiroplasma* of *M. achilles* is associated with the accumulation of new genes acquired through interactions with phages. Furthermore, the sAch, sAma and sRhe assemblies encode for six, four and five different plasmids respectively. They are all characterized by substantial higher level of read coverage than the genome contigs (Figure 1) suggesting the presence of multiple identical copies per bacterial cells. Interestingly, we failed to identify in our *Morpho Spiroplasma* genomes any CRISPR locus that are known to confer anti-viral protection in prokaryotes.

### Toxin genes identified in the Spiroplasma genomes of healthy Morpho males

As insects *Spiroplasma* are known to induce striking phenotypes in their host, such as male-killing promoted by the Spaid toxin or protection against parasites controlled by RIP-like enzymes (Harumoto and Lemaitre 2018; Ballinger and Perlman 2019), we specifically searched for the corresponding genes. One of the plasmids detected in both the sAch and sAma assemblies encodes for an ORF that combines both a RIP locus and a complete and structurally conserved Spaid gene (Figure 4B). This apparent bi-functional gene encodes for two RIP proteins in the 5’ end and a Spaid protein in 3’ end. The latter includes both ankyrin repeats (Ank) and a deubiquitinase domain (OTU), which are known to occur in the *Spiroplasma* strain sMel (Figure 4B) and induce male-killing in *Drosophila melanogaster* embryo. We observed ankyrin repeats, the OTU domain, and RIP domains in other genomes of *Spiroplasma* (Figure 4A). In particular, four homologous copies of RIP/OTU/Ank domains were also present in the sRhe assembly, and all of them are located on four different contigs. Although RIP-encoding genes are present in various *Spiroplasma* genomes (Figure 4A), the fusion of the RIP domain with the Spaid domain is an original feature found only in all of the *Morpho Spiroplasma* genomes as well as in a *Spiroplasma* found in the Hesperiid *Jemadia suekentonmiller (sSue)* (Moore & Ballinger 2023)(Figure 4B). Interestingly, *Spirosplama* associated with *Morpho* and *Jemadia* butterflies are phylogenetically closely related (Supplementary Figure 7) suggesting a common origin of the gene fusion rather than independent events. These features open the possibility that *Spiroplasma* endosymbionts may induce some peculiar phenotype in their South American butterfly hosts.

**Figure 4:**
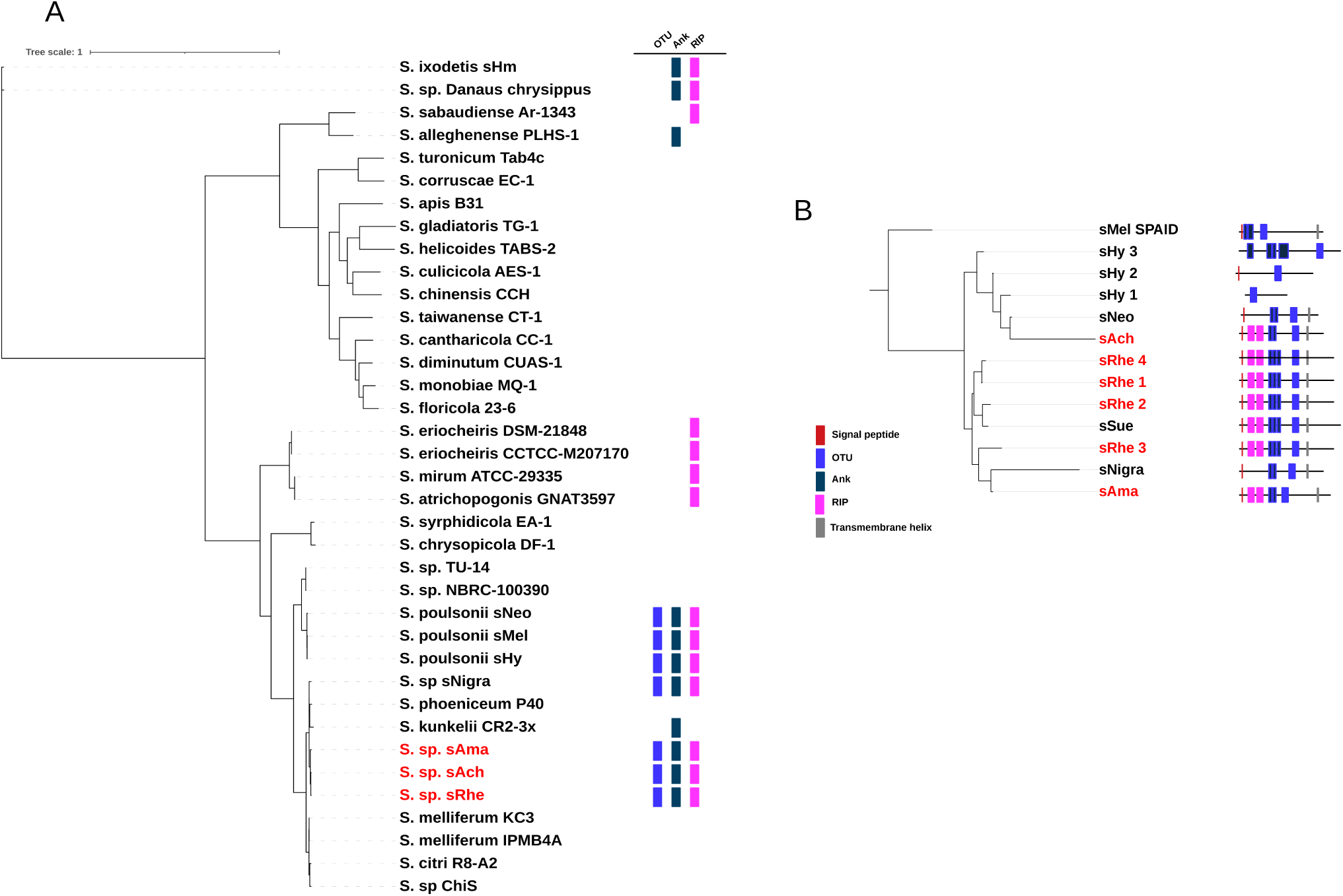
Distribution and organization of the toxin genes found in genomes of *Spiroplasma*. **A**: Distribution of the OTU (blue), Ankyrin (grey) and RIP (fucsia) encoding domains across the *Spiroplasma* whole-genome phylogeny. **B**. Phylogeny based on the OTU domain alignment of the Spaid-like proteins. Domain prediction based on Pfam similarity with known domains. The position of the *Morpho Spiroplasma* is highlighted in red in both trees.

### Horizontal and vertical transfer of Spiroplasma in Morpho

To estimate the prevalence of *Spiroplasma* within species of *Morpho* and test for horizontal *vs*. vertical transfer of this endosymbionts, we searched for the presence of *Spiroplasma* in different populations of *M. achilles* and its sister and sympatric species *M. helenor* (see Figure 5). We detected genomes of *Spiroplasma* with assembly size >100 kb and highly matching the sAch assembly in 26 out of 33 individuals of *M. achilles*; only 6 individuals had few contigs matching sAch (with assembly size <100kb), and a single one lacked any genomic trace of it (Figure 5 and Supplementary Table 1).

**Figure 5:**
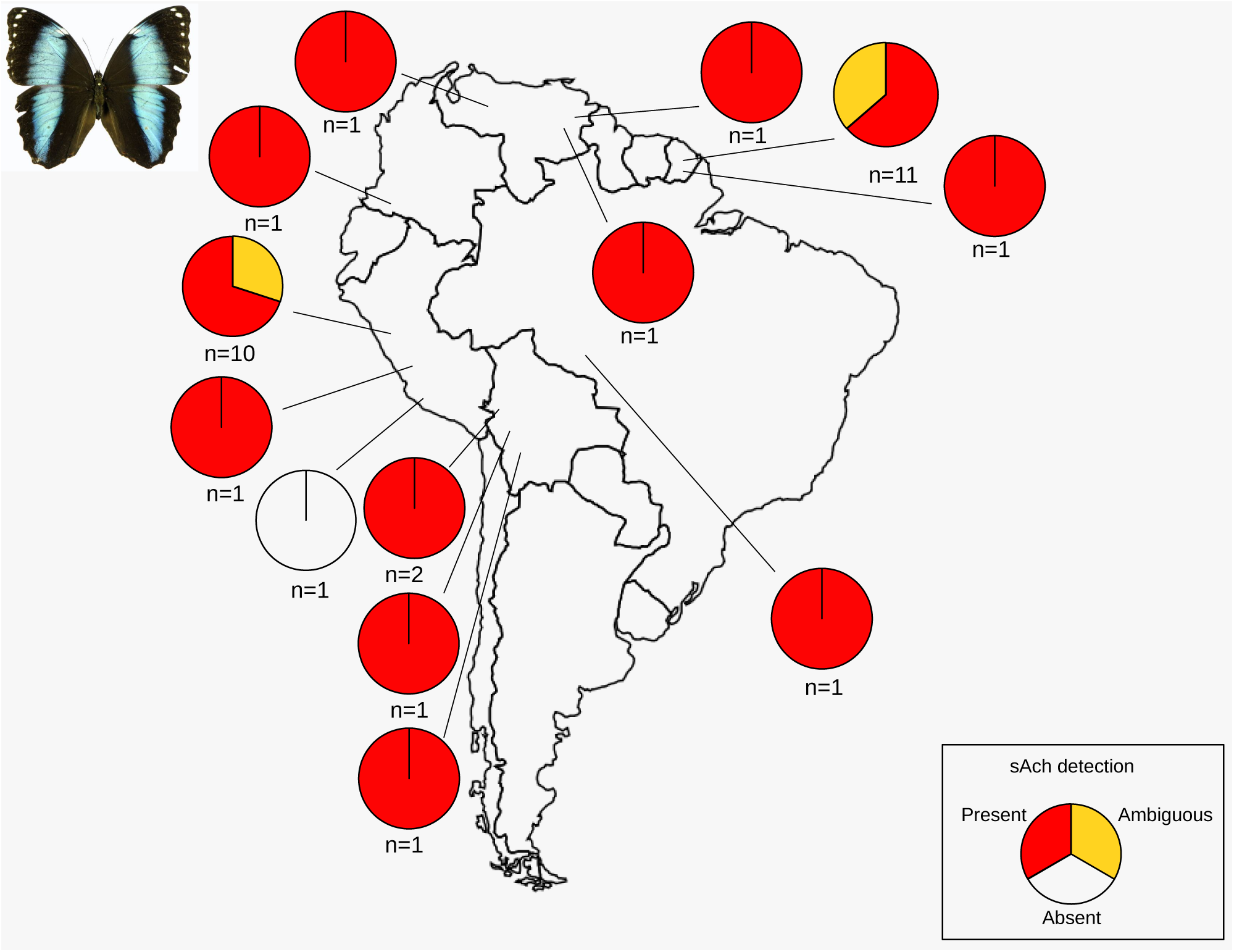
Geographic distribution of *Spiroplasma* in populations of *Morpho achilles*. Each circle represents a population of *M. achilles* and the number of individuals sampled per population is indicated on the bottom. The presence of *Spiroplasma* in each population is color coded as red (presence), yellow (ambiguous) and white (absence). See also Supplementary Figure 9.

Thus, all populations of *M. achilles* across south-America had *Spiroplasma*, except for one population in Peru represented by a single individual in our study (Figure 5). By contrast, among the 43 *M. helenor* individuals from the 27 populations we investigated, only a single individual had *Spiroplasma*. Therefore, although *M. achilles* and *M helenor* are sympatric species throughout the Amazonian basin and are closely-related species (3,6 million years of divergence, see Chazot et al. 2021), they display completely opposite patterns of infection by *Spiroplasma*.

Most of the *Spiroplasma* identified here had large genome assemblies (1 - 4 Mb) albeit the use of short-read sequencing technology that generally failed to assemble highly repeated regions. Moreover, some assemblies reached sizes comparable to the reference sAch assembly (Supplementary Figure 9). Homologous RIP and Spaid toxin genes were also found almost universally (Supplementary Figure 9).

To gain insight into the mode of transmission of the *Spiroplasma* symbionts in *Morpho* we performed reconciliation analysis between the symbiont phylogeny (whole mitochondrial phylogeny) and the host phylogeny (133 conserved *Spiroplasma* genes) (Figure 6A for the species dataset analysis, Figure 6B for the population dataset analysis and Supplementary Figure 10 for the combined global analysis). In the host tree, *M achilles* sequences form a monophyletic group with *M. helenor* as a sister group and *M. rhetenor* and *M. amathonte* form a monophyletic group that appears as a sister taxa of the *M achilles/M. helenor* clade (Figure 6A). By contrast, in the symbiont phylogeny, the unique *Spiroplasma* strain retrieved from *M. helenor* (sHel) and the sRhe *Spiroplasma* from *M. rhetenor* both appear well nested into the *M. achilles* clade (Figure 6A). This incongruence indicates a putative horizontal transmission of *Spiroplasma* between *M. helenor*, *M. achilles* and *M. rhetenor,* both living in sympatry in their range of distribution. By contrast, the tree of the *Spiroplasma* infecting *M. achilles* populations and the *M. achilles* host tree are congruent suggesting that both the endosymbiont and mtDNA are maternally inherited in *M. achilles* (Figure 6B). Thus, our findings strongly suggests a lateral exchange of *Spiroplasma* between *M. achilles* and the unique *M. helenor* individual carrying the symbiont. A possible host switch has also been evidenced between *M. achilles* and *M. rhetenor* albeit we lack population genomic data for the latter species to strengthen this result. On the other hand, the high congruence between the *Spiroplasma* and the mitochondrial tree within *M. achilles* agrees with a predominant vertical maternal transmission.

**Figure 6:**
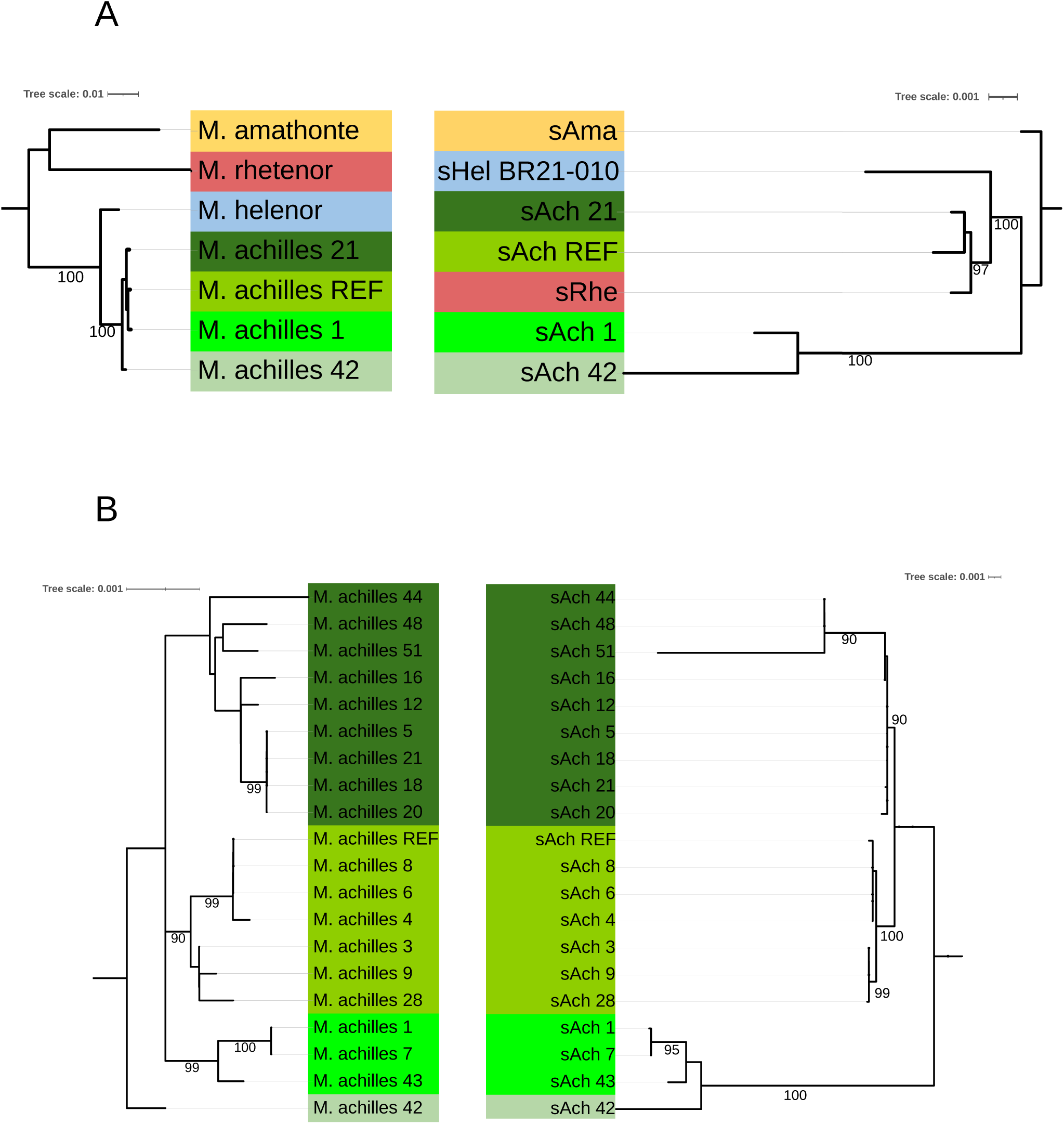
Reconciliation analysis between the host tree contrasted against the endosymbiont *Spiroplasma* tree to identify host lateral switches. (A) Genus-level analysis: The whole mitochondrial genome tree for the different *Morpho* species are compared with a 133 genes tree for *Spiroplasma.* (B) Population-level analysis: The mitochondrial genome tree of *M. achilles* populations is contrasted with a 133 genes tree for the corresponding *Spiroplasma* symbionts. Ultra-fast boostraps values are indicated on each branch. Color blocs correspond to the main phylogenetic clusters identified in the species phylogeny. The complete phylogenies with the prediction of the possible horizontal host switches are available in Supplementary Figure 10.

## Discussion

### Peculiar evolution of Spiroplasma genomes in Morpho butterflies

We documented the presence of the bacterial endosymbiont *Spiroplasma* in four out of 11 species of *Morpho* butterflies studied here, highlighting that the association with this endosymbiont is likely to greatly vary across closely related host species, even when they live in sympatry. *Spiroplasma* retrieved from *Morpho* belongs to the Citri clade whereas *Spiroplasma* symbiont infecting *Danaus* butterflies belong to the distantly related *Ixodetis* clade. Interestingly, *Spiroplasma* strains infecting hesperiid and nymphalid butterflies that also belong to the Citri clade have been recently reported, including in species found in South America (Moore & Ballinger 2023, Twort et al. 2022). Although nymphalid and hesperiid are distantly related butterflies, *Spiroplasma* infecting the hesperiid *Jemadia suekentonmiller* appears very closely related to the *Spiroplasma* associated with *Morpho* nymphalid butterflies (Supplementary Figure 7). These results strongly suggest symbiont horizontal transfer among distantly-related host species. The phylogenetic discrepancies between *Spiroplasma* genes and mitochondrial genes across *Morpho* species also suggests that horizontal transfer between species living in sympatry might occur. The circulation of *Spiroplasma* in the hemolymph is thought to facilitate horizontal transfer across sympatric species, for instance through the consumption of hemolymph by mites (Jaenike et al. 2010). Because the *Spiroplasma* found in *Morpho* butterflies is closely related to a strain documented as a plant pathogen, the inter-specific transmission of these symbionts could also be enabled by the shared consumption of host plants by caterpillars. The three *Morpho* species presenting close *Spiroplasma* genomes (*M. achilles*, *M. helenor* and *M. rhetenor)* share several hostplants in French Guyana, supporting this hypothesis (Anon 2017). The presence of *Spiroplasma* in *Morpho* butterflies might thus stem from interactions with other arthropods or through host plant consumption. However, population genomics of *Spiroplasma* within the species *M. achilles* species also indicate widespread distribution and predominance of vertical transmission within species.

Interestingly, our results highlight the occurrence of *Spiroplasma* symbiont in *M. amathonte* and *M. achilles*, two species that have diverged from more than 17 My (Chazot et al. 2021) and that currently occupying non-overlapping geographical ranges: *M. amathonte* is found in central America and in the western slopes of the Andes while *M. achilles* inhabits the Amazonian basin. The weak genome synteny of the *Spiroplasma* genomes of these two species could result from independent genomic evolution of the *Spiroplasma* strains in these two hosts. Moreover, the congruence of host and symbiont phylogenies support a vertical transmission from a common ancestor, followed by independent evolution within each host species. Thus, the ubiquitous presence of *Spiroplasma* in *M. achilles* populations across south-America and the predominant vertical transmission of the symbiont at various evolutionary time-scale points at a stable association between *Spiroplasma* symbiont and *Morpho* butterflies. The lack of detection of *Spiroplasma* in some *Morpho* species (e.g. *M. menelaus*, closely related to *M. amathonte*) would then suggest a secondary loss, although the sequencing of a single individual per species does not allow to robustly conclude on the absence of *Spiroplasma* in some *Morpho* species.

### Massive genome size promoted by large expansion in diverse mobile genetic elements

While endosymbionts as *Spiroplasma* display streamlined genomes (Gerth et al. 2021), the genome size of the *Spiroplasma* observed in our study is surprisingly large compared to other *Spiroplasma* genome sequenced to date (from 0,9 Mb to 1,9 Mb, average = 1,2Mb). This result is unlikely to be caused by the quality of our assembly relaying on long-read sequencing technology: 72% of the *Spiroplasma* genome infecting *M. achilles* used in our comparative analysis correspond to a single contig genome assembly. Moreover, the use of different assembly softwares both leads to large assemblies (>4Mb). The assemblies of other *Spiroplasma* used here for comparisons have been selected to have comparable genome metrics (contig N50 > 100 kb) to discard highly fragmented assemblies that may capture only a fraction of the actual *Spiroplasma* genomes. In addition, the extremely low GC% of the *Spiroplasma* genomes compared to host genomes ease the binning process to discriminate the contigs that belong to the host and those corresponding to the symbionts. In addition, none of the ORFs encoded by the different *Spiroplasma* assemblies have their best matches with any eukaryotic sequences, strongly arguing that these contigs do not correspond to integrated fragments in the host genome. As *Spiroplasma* also infects plants, the detection of *Spiroplasma* in our *Morpho* assemblies might have been the result of contamination with plant DNA ingested by the butterflies. However, virtually none of the eukaryotic contigs binned in our analyses matches with plant: only seven small contigs in the *M. achilles* genome assembly match with upland cotton sequences (*Gossypium hirsutum*) which are not known to be a *Morpho* feeding source. Thus, the lack of plant contigs in our assemblies indicated that DNA contamination with *Spiroplasma* infecting plant is unlikely.

Although *Morpho Spiroplasma* genomes are much bigger than other *Spiroplasma* genomes, they encode for a similar number of orthologous gene groups compared to other *Spiroplasma* (around 1000 groups of orthology). Our analyses indicate that recent and massive expansion of diverse mobile genetic elements (MGE) are responsible for this striking inflation in genome size. Record-level of Insertion Sequences (accounting for 1550 kb), integrated prophages (538kb) and plasmids (278 kb) represent nearly 60% of the total genome size of the *Spiroplasma* found in *M. achilles* (accounting for 2366 kb on a total genome size of 4075 kb). All *Spiroplasma* genomes detected in *Morpho* butterflies display such extreme expansion of MGE, in contrast with the relative paucity of MGE generally found in *Spiroplasma* (Gerth et al. 2021) or *Wolbachia* genomes (Cerveau et al. 2011). In *Spiroplasma*, it has been swown that strains harbouring CRISPR/Cas anti-viral systems display low level of prophage density (Gerth et al. 2021). On the other hand, strains for which the CRISPR/Cas have been lost display multiple prophage sequences. The absence of CRISPR loci in our *Morpho Spiroplasma* assemblies fit with this model and suggests that our *Spiroplasma* symbionts have lost the capacity to control phage infection and prophage insertions. Moreover, most of the IS copies found in *Spiroplasma* of *Morpho* butterflies are recently-transposed elements, indicating an ongoing and continuous accumulation. Such MGE proliferation has also been sporadically observed in *Orientia* symbionts (Bacteria: Rickettsiales), a widespread *Rickettsia*- like, intra-cellular bacteria associated with mites (Batty et al. 2018) and in *Mycoplasma* endosymbionts, associated with diverse fungi (Naito and Pawlowska 2016). In addition, integrated phage genomes provide numerous new genes and functions in *Spiroplasma* associated with *Morpho* butterflies. The massive expansion of MGEs leads to an inflated genome size but also provides a source of new genes and functions expanding the diversity of the genomic repertoire of *Spiroplasma* symbionts infecting *Morpho* butterflies. We propose that the recombinations induced by MGEs and the gene flow provided by phage genome integration have counter-balanced the genome erosion process in the S*piroplasma* of *Morpho*, which contrasts with the important genome size reduction observed in most endosymbionts. In this regard, it’s noteworthy that a very similar scenario has been recently been evidenced in *Arsenophonus* symbionts in which vertically inherited strains has larger genome than horizontally transmitting symbionts (Siozios et al. 2024). As for *Morpho Spiroplasma*, resistance to genome erosion in *Arsenophonus* is also linked to MGE expansion and loss of CRISPR-cas defense systems. Such rapid evolution might stem from peculiar adaptation in the symbionts of *Morpho*, allowing stable association and high prevalence in some *Morpho* species. For instance, MGE-encoded toxin genes might have contributed to increase the symbiont persistence in some *Morpho* butterfly populations.

### Evolution of toxin genes and putative protective effect

By detecting *Spiroplasma* in adult *Morpho* males sampled in the wild, our study suggests that the presence of *Spiroplasma* does not prevent the development of males in *Morpho* butterflies, in sharp discrepancy with the male-killing effects reported in the butterfly *Danaus chrysippus* (Jiggins et al. 2000), but similar to the results obtained with *Wolbachia* in Neotropical Acraeini (Nymphalidae) (Silva-Brandão et al. 2021). More specifically, the toxin gene *Spaid* found in *Spiroplasma poulsonii* and documented to trigger male killing in *Drosophila* (Harumoto and Lemaitre 2018) does not have the male-killing (MK) effect in *Morpho*. Since *Spaid* sequences observed in *Morpho* and *Drosophila* are largely divergent, the evolution the *Spaid* gene might be associated with changes in gene function. Alternatively, resistance to the sex-ratio distortion effect of the *Spiroplasma Spaid* toxins may thus have evolved in *Morpho* butterflies. Such MK-suppression have indeed been observed in plant-hopper (Yoshida et al. 2021) and in lacewing (Hayashi et al. 2018). Alternatively, as *Spaid* toxins target the gene dosage compensation system that increases the transcription of genes on the male single X chromosome in *Drosophila* (Harumoto & Lemaitre 2018), this toxin could thus be ineffective on the ZW sex-determination system where females are heterogametic. Supporting this view, MK-inducing *Spiroplasma* in Lepidoptera lack the *Spaid* toxins genes, but the genetic determinant(s) of the MK phenotype are unknown (Arai et al. 2022). However, strong conservation of the *Spaid* genes among *Morpho* species and populations could be consistent with an associated selective advantage.

Our study also reveals the evolution of the specific architecture of toxin genes in the *Spiroplasma* of *Morpho* butterflies, including *RIP* physical linkage with the *Spaid* gene. Among *Spiroplasma*, this original gene structure has only been reported in the neotropical skipper butterfly *Jemadia suekentonmiller* (Hesperiidae: Pyrrhopyginae) (Moore & Ballinger 2023). While the functional implication of this evolution cannot be inferred from our current results, the localization of these genes on plasmids also suggests that they may spread across bacteria. Their persistence might have been promoted by natural selection, either because they act as selfish elements or because of positive impact on host fitness. In *Drosophila*, RIP proteins produced by *Spiroplasma poulsonii* have indeed been documented to induce positive effects on host survival, through their protective effect (Moore & Ballinger 2023). Such defensive effect of *Spiroplasma* could have a positive impact on the fitness of *Morpho* butterflies and might explain their high prevalence in *M. achilles*.

### Contrasted prevalence of Spiroplasma in sympatric sister species

The evolution of the *Spaid* gene might have resulted in a change of function in *Morpho* butterflies, disabling the male-killing mechanism. Alternatively, resistance to male-killing effect might have evolved in *Morpho* butterflies. The evolution of MK-suppression has been documented in natural populations of the butterfly *Hypolimnas bolina* (Nymphalidae) infected by *Wolbachia* (Hornett et al. 2022). Such evolution of resistance is likely to be under strong positive selection given the high fitness costs for the hosts induced by male-killing genes (Hornett et al. 2022). The *Spiroplasma* is highly prevalent in *M. achilles* and quite rare in the sympatric sister species *M. helenor*. This might suggest that resistance to the deleterious effect of *Spiroplasma* could be restricted to *M. achilles*.

Cytoplasmic incompatibilities induced by the symbiont might explain the huge difference in its prevalence between these two sister-species living in sympatry. *Spiroplasma*-induced cytoplasmic incompatibilities have been recently documented in the wasp *Lariophagus distinguendus* (Pollman et al. 2022). Such incompatibilities might explain the difference in the *Spiroplasma* prevalence between the two sister-species *M. achilles* and *M. helenor* living in sympatry. In case of CI, crosses between infected males and uninfected females generally do not produce offspring; CI could thus limit genetic exchange between the two sympatric species. Altogether, our current results on the contrasted *Spiroplasma* prevalence in these sister-species therefore raises the question of the potential impact of this endosymbiont on barrier to gene flow between these sympatric species. Endosymbionts like *Spiroplasma* and *Wolbachia* have indeed been suggested to generate post-zygotic barriers to gene flow (see Duplouy & Hornett 2018 for a review), but their role in initiating vs. reinforcing speciation remains largely uncovered. Our study therefore points at the needs to investigate the effect of *Spiroplasma* on reproductive isolation and its significance for species diversification and co- existence in sympatry.

## Conclusion

Studies on heritable symbionts in natural populations generally support a dynamic model of gene gain and loss shaped by the ecological interactions with their host. On the other hand, heritable symbiont lifestyle induces genetic isolation and population bottlenecks that lead to mutational decay and genome streamlining. In comparison with *Spiroplasma* genomes retrieved from other insects, *Morpho Spiroplasma* genomes display a massive expansion of diverse mobile genetic elements as transposable elements, prophages or plasmids. In addition, we documented a peculiar arrangements of toxin RIP and Spaid-encoding genes in the *Spiroplasma* of *Morpho* species, that might enhance protection of the butterflies against parasites. The study of *Spiroplasma* symbionts in natural population of diverse *Morpho* butterfly species supports a stable association in *Morpho achilles* populations across south-America, whereas *Spiroplasma* appears very rare in sympatric *M. helenor* populations. This contrasted symbiont distribution among sympatric *Morpho* species is associated with a global predominant vertical transmission of the symbiont. *Morpho Spiroplasma* genomes also display a remarkable resistance to genome erosion by the mean of massive expansions of diverse mobile genetic elements as transposable elements, prophages or plasmids. In addition, the co-localisation of the toxin RIP and Spaid-encoding genes might be the key-driver of this lasting association by conferring host protection against parasite. Our study calls for additional investigations of the phenotypic effects of the *Spiroplasma* on their butterfly hosts to better understand how their ecological interactions shapes – and is shaped by – their evolution.

## Acknowledgments

We would like to thank Heloïse Bastide and David Ogereau for their helps at the initial phase of this work, Josephine Ledamoisel, Rémi Mauxion and Owen MacMillan for their support during fieldwork in Panama, as well as Mathieu Chouteau and Mélanie McClure during fieldwork in Peru and French Guiana. Some of the analyses and simulations were performed on the the Plateforme de Calcul Intensif et Algorithmique PCIA (MNHN/CNRS), on the MeSU platform at Sorbonne-Université and on the Genotoul bioinformatics platform Toulouse Occitanie (Bioinfo Genotoul). André V.L. Freitas thanks the Conselho Nacional de Desenvolvimento Científico e Tecnológico – CNPq (fellowship 304291/2020-0), and the FAPESP (grant 2021/03868-8). Thank you to the ICMBio for collecting permits (10438-2 to 10438-4). This study is registered in the Brazilian SISGEN (A43CEB1).

## Data Availability Statement

PacBio *(genus dataset*) and Illumina (*sister-species dataset*) read sequences and genome assemblies generated in this study were deposited in the NCBI database under the bioprojects PRJNA1069011 and PRJNA1063620. Primary raw data for each analysis can be downloaded at https://doi.org/10.6084/m9.figshare.24582867.v1

## Funding

This work was supported by the European Research Council (ERC) Consolidator Grant (ERC-2022-COG - OUTOFTHEBLUE – 101088089), ANR-tremplin and the CNRS MITI project ECODEMO. Views and opinions expressed are however those of the authors only and do not necessarily reflect those of the European Union or the European Research Council. Neither the European Union nor the granting authority can be held responsible for them.

## Notes

### Competing Interest Statement

The authors have declared no competing interest.

### Summary of Updates

Title, summary, some figures and the discussion section have been revised. Supplementary files updated.

