## Supplementary Figures and Table for "Genome evolution and between-host transmission of *Spiroplasma* endosymbiont in wild communities of Morpho butterflies"

### Supplementary Table 1 : *Morpho achilles* and *M. helenor* sample distribution and prevalence of *Spiroplasma* genome sequences and putative toxin genes

| Sample | Species | Location | Mapped reads (nt) | Assembly size (nt) | BLAST assignation | Spaid/RIP assembly size (nt) | RIP BLAST assignation | Spaid BLAST assignation |
|---|---|---|---|---|---|---|---|---|
| VL16-1009 | M. helenor | Urahuasha (Peru) | 14020366 | 3336 | Absent | 301 | Absent | Absent |
| VL16-1005 | M. helenor | Urahuasha (Peru) | 3696682 | 1467 | Absent | 0 | Absent | Absent |
| VE21-002 | M. helenor | San Franscico de Guayo, Orinoco delta (Venezuela) | 58387834 | 289540 | Absent | 567 | Absent | Absent |
| VE21-001 | M. helenor | Catetimbo zulia (Venezuela) | 41650223 | 162634 | Absent | 0 | Absent | Absent |
| TR21-001 | M. helenor | Tunaipuna (Trinidad y Tobago) | 72517101 | 428144 | ambiguous | 0 | Absent | Absent |
| PA21-001 | M. helenor | Colon (Panama) | 8542766 | 3354 | Absent | 0 | Absent | Absent |
| GU21-001 | M. helenor | Los Tarrales (Guatemala) | 82890077 | 467224 | Absent | 265 | Absent | Absent |
| FG19-058 | M. helenor | Cascade de Kaw (French Guiana) | 6586041 | 1973 | Absent | 557 | Absent | Absent |
| FG19-014 | M. helenor | Cascade de Kaw (French Guiana) | 6673236 | 2442 | Absent | 302 | Absent | Absent |
| CR21-003 | M. helenor | Kirby Woik Finca (Costa Rica) | 16064115 | 3738 | Absent | 279 | Absent | Absent |
| CR21-002 | M. helenor | San Vito (Costa Rica) | 102254925 | 532068 | Absent | 0 | Absent | Absent |
| CR21-001 | M. helenor | San Vito (Costa Rica) | 24848282 | 45539 | Absent | 0 | Absent | Absent |
| CO21-M5970 | M. helenor | Putomayo (Colombia) | 5376692 | 2627 | Absent | 575 | Absent | Absent |
| CO21-M5969 | M. helenor | Putomayo (Colombia) | 7013899 | 1786 | Absent | 576 | Absent | Absent |
| CO21-M5968 | M. helenor | Putomayo (Colombia) | 17350523 | 41345 | Absent | 1831 | Absent | Absent |
| CO21-M5961 | M. helenor | Boyaca (Colombia) | 13268945 | 6230 | Absent | 295 | Absent | Absent |
| CO21-M5959 | M. helenor | Boyaca (Colombia) | 9272746 | 4680 | Absent | 287 | Absent | Absent |
| CO21-M5958 | M. helenor | Boyaca (Colombia) | 11887150 | 2515 | Absent | 294 | Absent | Absent |
| CO21-M5957 | M. helenor | Boyaca (Colombia) | 9468373 | 2346 | Absent | 0 | Absent | Absent |
| CO21-M5911 | M. helenor | Valle del Cauca (Colombia) | 12516004 | 2856 | Absent | 0 | Absent | Absent |
| CO21-M5910 | M. helenor | Valle del Cauca (Colombia) | 12726546 | 384 | Absent | 0 | Absent | Absent |
| CO21-M5663 | M. helenor | Boyaca (Colombia) | 5424099 | 3261 | Absent | | Absent | Absent |
| CO21-M5625 | M. helenor | Boyaca (Colombia) | 12433570 | 4527 | Absent | 0 | Absent | Absent |
| CO21-M5624 | M. helenor | Boyaca (Colombia) | 5454654 | 2305 | Absent | 288 | Absent | Absent |
| BR21-023 | M. helenor | Jundiai (Brazil) | 6451817 | 2124 | Absent | 278 | Absent | Absent |
| BR21-022 | M. helenor | Intervales (Brazil) | 52880029 | 11395 | Absent | 0 | Absent | Absent |
| BR21-017 | M. helenor | Santarém (Brazil) | 1670384 | 381 | Absent | 0 | Absent | Absent |
| BR21-016 | M. helenor | Almeirim (Brazil) | 13097682 | 2172 | Absent | 285 | Absent | Absent |
| BR21-014_ | M. helenor | Porto Velho (Brazil) | 10829752 | 6679 | Absent | 559 | Absent | Absent |
| BR21-013 | M. helenor | Porto Velho (Brazil) | 8407917 | 7365 | Absent | 581 | Absent | Absent |
| BR21-011 | M. helenor | Querência (Brazil) | 10042572 | 3730 | Absent | 0 | Absent | Absent |
| BR21-010 | M. helenor | Tapajós (Brazil) | 62896131 | 993039 | Present | 39440 | Present | Present |

| | | | | | | | | |
|---|---|---|---|---|---|---|---|---|
| BR21-009 | M. helenor | Tapajós (Brazil) | 57443680 | 9385 | Absent | 0 | Absent | Absent |
| BR21-008 | M. helenor | Vitória da conquista (Brazil) | 11857494 | 3800 | Absent | 267 | Absent | Absent |
| BR21-006 | M. helenor | Vicosa (Brazil) | 5922799 | 747 | Absent | 299 | Absent | Absent |
| BR21-005 | M. helenor | Serra Grande (Brazil) | 6318979 | 1923 | Absent | 600 | Absent | Absent |
| BR21-004 | M. helenor | Atibaia (Brazil) | 11457884 | 4921 | Absent | 286 | Absent | Absent |
| BR21-003 | M. helenor | Porto Ferreira | 22068535 | 32182 | Absent | 535 | Absent | Absent |
| BR21-001 | M. helenor | Aiuruoca (Brazil), | 10239720 | 6363 | Absent | 568 | Absent | Absent |
| BO21-001 | M. helenor | Bolivia Muyupampa | 8663859 | 6802 | Absent | 279 | Absent | Absent |
| 9 | M. achilles | Cascade de Kaw (French Guiana) | 407875746 | 1196252 | Present | 99408 | Present | Present |
| 8 | M. achilles | Cascade de Kaw (French Guiana) | 247582483 | 1247416 | Present | 111233 | Present | Present |
| 7 | M. achilles | Cascade de Kaw (French Guiana) | 285666797 | 995976 | Present | 39944 | Present | Present |
| 6 | M. achilles | Cascade de Kaw (French Guiana) | 132787974 | 1158694 | Present | 96362 | Absent | Present |
| 51 | M. achilles | Rio Undumo (Bolivia) | 181992682 | 1745805 | Present | 86439 | Present | Present |
| 50 | M. achilles | Caranavi (Bolivia) | 184789866 | 1319148 | Present | 1136 | Present | Present |
| 5 | M. achilles | Cascade de Kaw (French Guiana) | 14558846 | 9176 | ambiguous | 8364 | Absent | Absent |
| 48 | M. achilles | Rio Ene (Perou, Central) | 269387385 | 1592410 | Present | 98104 | Present | Present |
| 47 | M. achilles | Caranavi (Bolivia) | 241378677 | 1773081 | Present | 844 | Absent | Absent |
| 46 | M. achilles | Rio Caripe (Bolivia) | 155250672 | 784370 | Present | 906 | Absent | Absent |
| 45 | M. achilles | Bolivar (venezuela) | 424174736 | 3012551 | Present | 9940 | Absent | Absent |
| 44 | M. achilles | Bauxilum (Venezuela ouest) | 369729120 | 2009995 | Present | 129538 | Present | Present |
| 43 | M. achilles | Mocoa Colombie) | 294829446 | 1966354 | Present | 58444 | Absent | Present |
| 42 | M. achilles | Rondônia (Brésil) | 580603055 | 4037608 | Present | 60855 | Absent | Absent |
| 41 | M. achilles | Rio Grande (Venezueal) | 577758966 | 3206874 | Present | 867 | Absent | Absent |
| 4 | M. achilles | Cascade de Kaw (French Guiana) | 231551027 | 1182833 | Present | 107199 | Present | Present |
| 31 | M. achilles | Cascade de Kaw (French Guiana) | 20347379 | 5183 | ambiguous | 842 | Absent | Absent |
| 30 | M. achilles | Cascade de Kaw (French Guiana) | 27450108 | 52141 | ambiguous | 6578 | Absent | Absent |
| 3 | M. achilles | Cascade de Kaw (French Guiana) | 266612469 | 1209569 | Present | 106264 | Present | Present |
| 28 | M. achilles | Cascade de Kaw (French Guiana) | 420620143 | 1237646 | Present | 117278 | Present | Present |
| 26 | M. helenor | Santa Fe (Panama) | 22119614 | 3152 | Absent | 525 | Absent | Absent |
| 25 | M. helenor | Campana (Panama) | 14600660 | 5660 | Absent | 0 | Absent | Absent |
| 24 | M. helenor | Santa Fe (Rio Mulaba) | 28190318 | 6412 | Absent | 0 | Absent | Absent |
| 22 | M. achilles | Alto_Shilcayo (Peru) | 18086584 | 11792 | ambiguous | 1152 | Absent | Absent |
| 21 | M. achilles | Alto_Shilcayo (Peru) | 116052913 | 1128408 | Present | 108345 | Present | Present |
| 20 | M. achilles | Alto_Shilcayo (Peru) | 185111407 | 1094483 | Present | 104789 | Present | Present |
| 2 | M. achilles | Cascade de Kaw (French Guiana) | 17817425 | 300681 | Present | 11619 | Present | Present |
| 19 | M. achilles | Alto_Shilcayo (Peru) | 21799435 | 9196 | ambiguous | 603 | Absent | Absent |
| 18 | M. achilles | Alto_Shilcayo (Peru) | 147144461 | 1105477 | Present | 93648 | Present | Present |

| | | | | | | | | |
|---|---|---|---|---|---|---|---|---|
| 16 | M. achilles | Alto_Shilcayo (Peru) | 81149907 | 1081244 | Present | 99415 | Present | Present |
| 15 | M. achilles | Alto_Shilcayo (Peru) | 150536398 | 1094387 | Present | 100351 | Present | Present |
| 14 | M. achilles | Alto_Shilcayo (Peru) | 7309755 | 1968 | ambiguous | 587 | Absent | Absent |
| 13 | M. achilles | Alto_Shilcayo (Peru) | 123187404 | 396512 | Present | 70519 | Present | Present |
| 12 | M. achilles | Alto_Shilcayo (Peru) | 78818582 | 1116188 | Present | 91822 | Present | Present |
| 11 | M. achilles | San Antonio (Peru) | 18803408 | 2413 | Absent | 566 | Absent | Absent |
| 1 | M. achilles | Piste du haut (French Guiana) | 377678547 | 1066629 | Present | 41285 | Absent | Present |

**Supplementary Table 2 : Characteristics of *Spiroplasma* assemblies using different genome assemblers**

| Dataset | Assembler | Assembly size (kb) | Contigs (#) | N50 (kb) | Completness (%) |
|---|---|---|---|---|---|
| *M. achilles* | Hifiasm | 4075 | 46 | 2667 | 98 |
| | Flye | 3011 | 22 | 167 | 56 |
| | HiCanu | Not Found | | | |
| *M. amathonte* | Hifiasm | 3144 | 16 | 2570 | 60 |
| | Flye | 4148 | 29 | 342 | 99 |
| | HiCanu | Not Found | | | |
| *M. rhetenor* | Hifiasm | 2860 | 44 | 157 | 78 |
| | Flye | 1186 | 28 | 114 | 35 |
| | HiCanu | Not Found | | | |

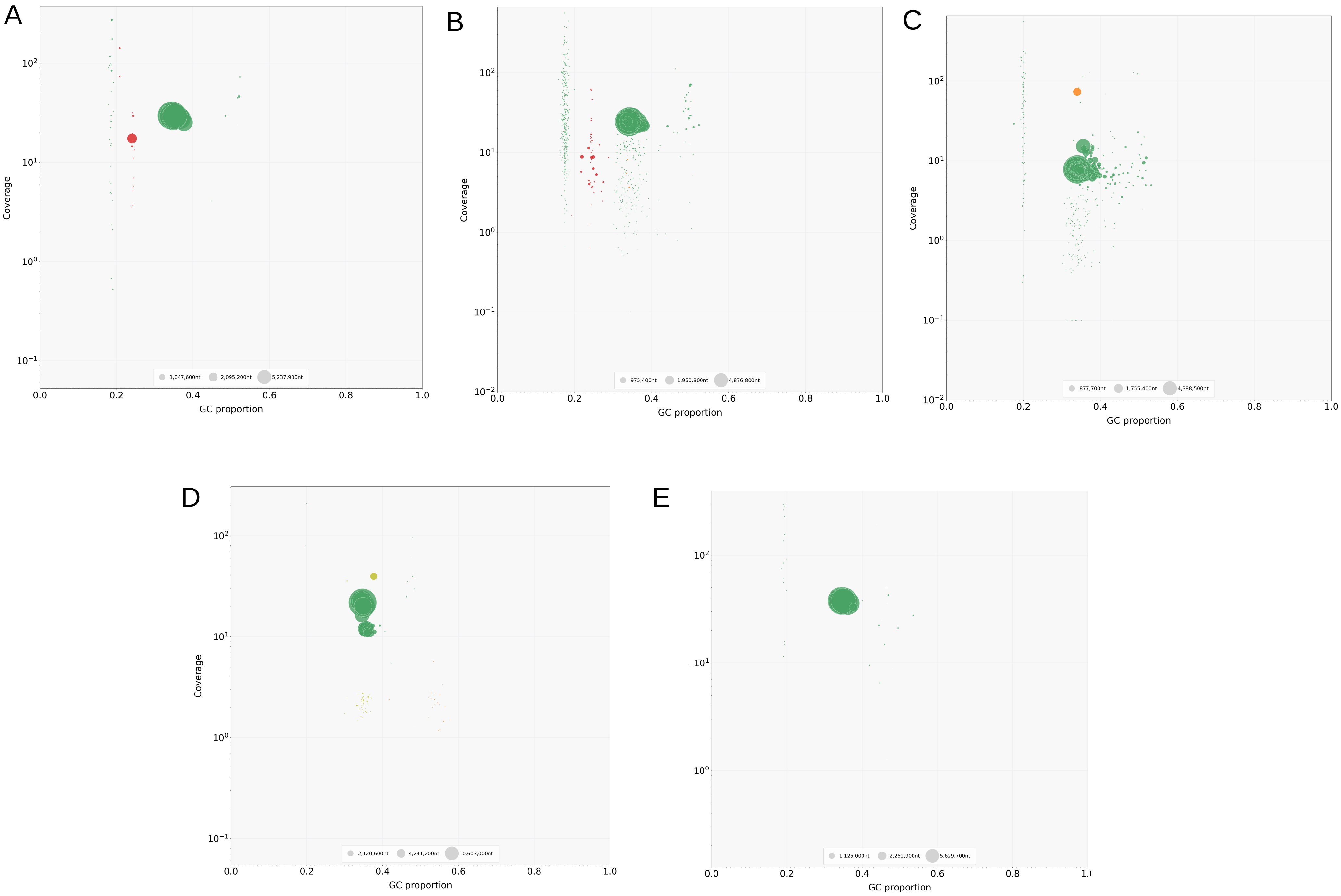
A
Coverage
GC proportion
1,047,600nt
2,095,200nt
5,237,900nt
B
Coverage
GC proportion
975,400nt
1,950,800nt
4,876,800nt
C
Coverage
GC proportion
877,700nt
1,755,400nt
4,388,500nt
D
Coverage
GC proportion
2,120,600nt
4,241,200nt
10,603,000nt
E
GC proportion
1,126,000nt
2,251,900nt
5,629,700nt

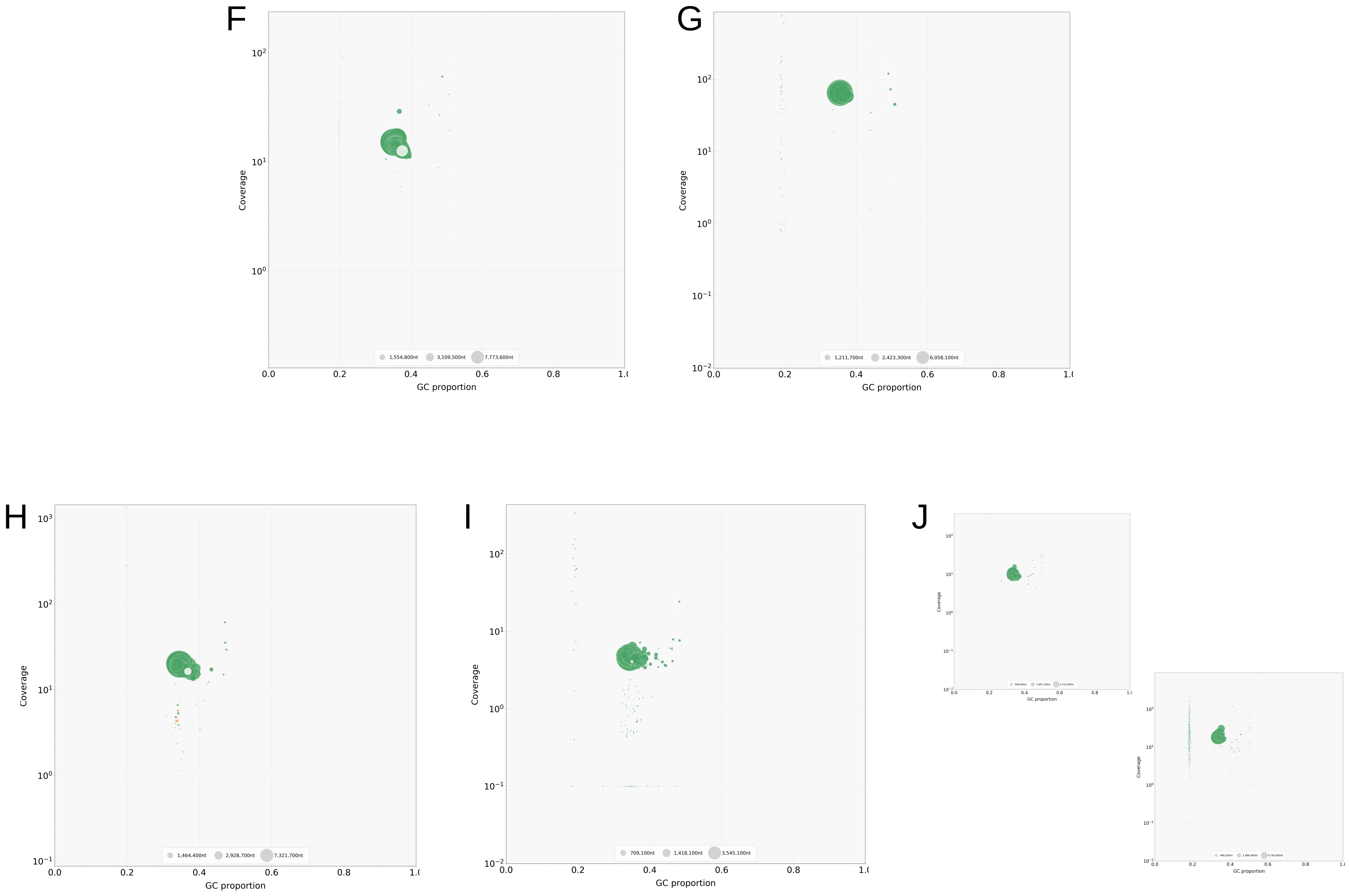


**Supplementary Figure 1: Blobtools of *Morpho* butterflies showing the presence of symbiont genomes in the assemblies: *M. amathonte* (A), *M. rhetenor* (B), *M. hecuba* (C), *M. helenor* (D), *M. marcus* (E), *M. deidamia* (F), *M. eugenia* (G), *M. granadensis* (H), *M. menelaus* (I) and *M. telemachus* gold (upper) and blue (lower)(J).** Contigs represented as circles were binned based on their GC%, read coverage, and taxonomic assignment. Dark green contigs matched arthropod sequences, red contigs matched Mollicutes (*Spiroplasma)*, orange contigs matched Proteobacteria (*Wolbachia*), and light green contigs matched Firmicutes (*Enterococcus*). The size of the circle is proportional to the size of the contigs.

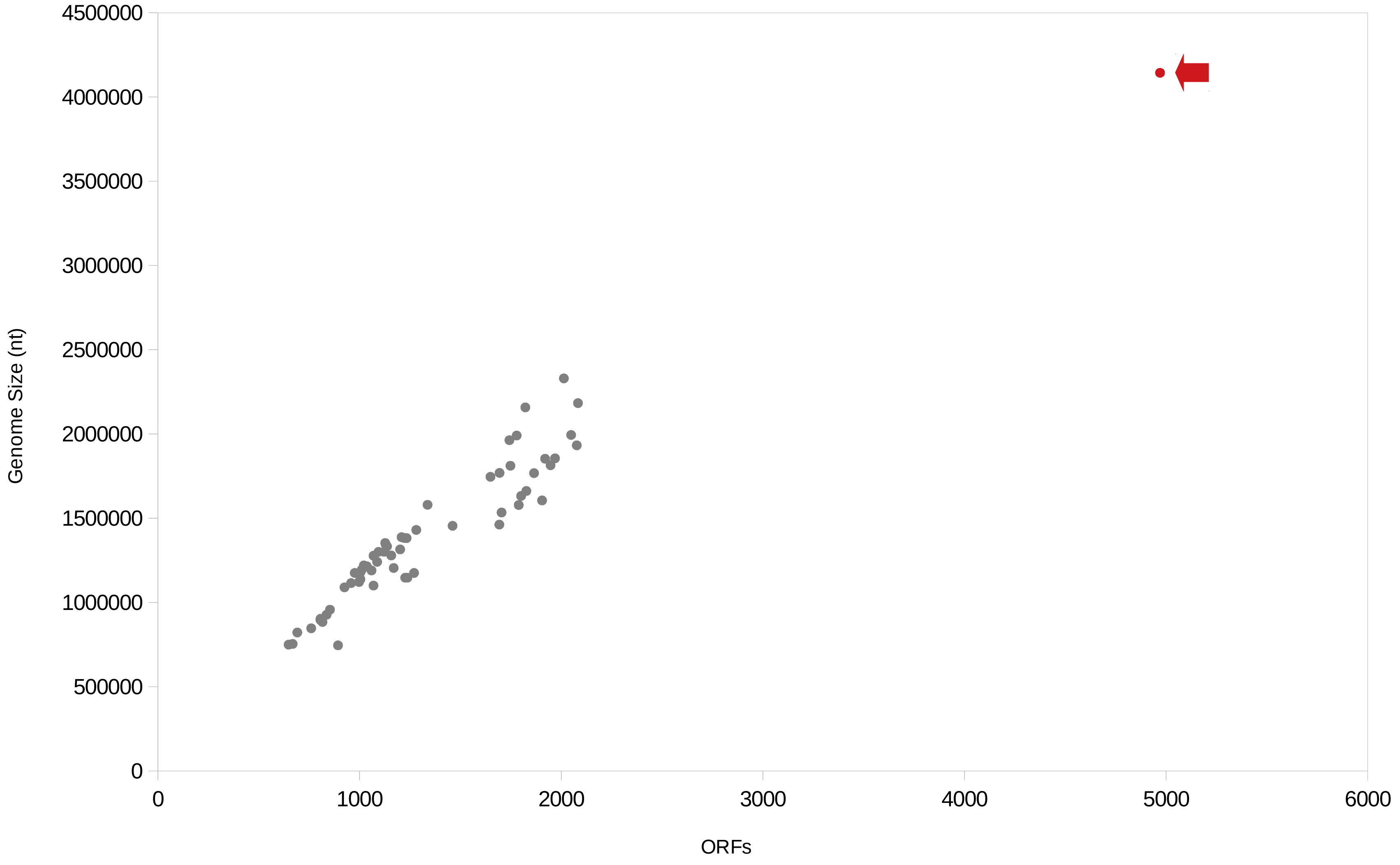


**Supplementary Figure 2: Number of ORFs of the 62 *Spiroplasma* genomes (gray dots) and *Morpho Spiroplasma* sAch (red dot with a red arrow) ploted with respect to their genome sizes.**

A.

B.

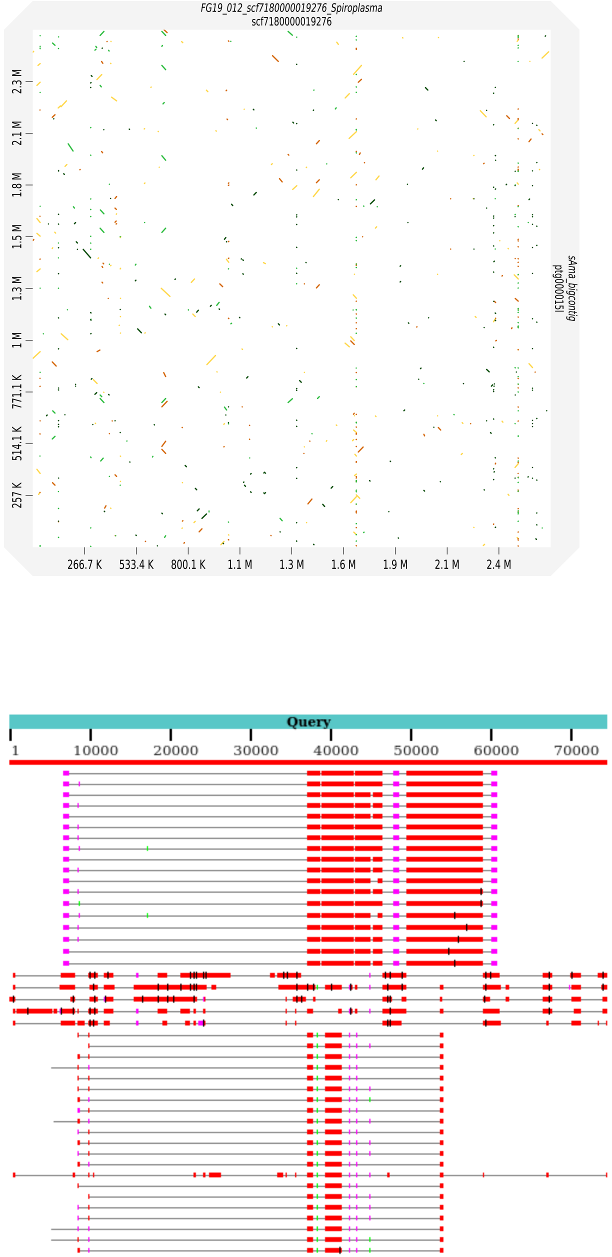


**Supplementary Figure 3: Alignement of the *Spiroplasma* genome assemblies.** Whole genome alignment of the sAch and the sAma genome assemblies (**A**) and alignement of the 46 small repetitive contigs present in the sAch genome assembly (**B**).

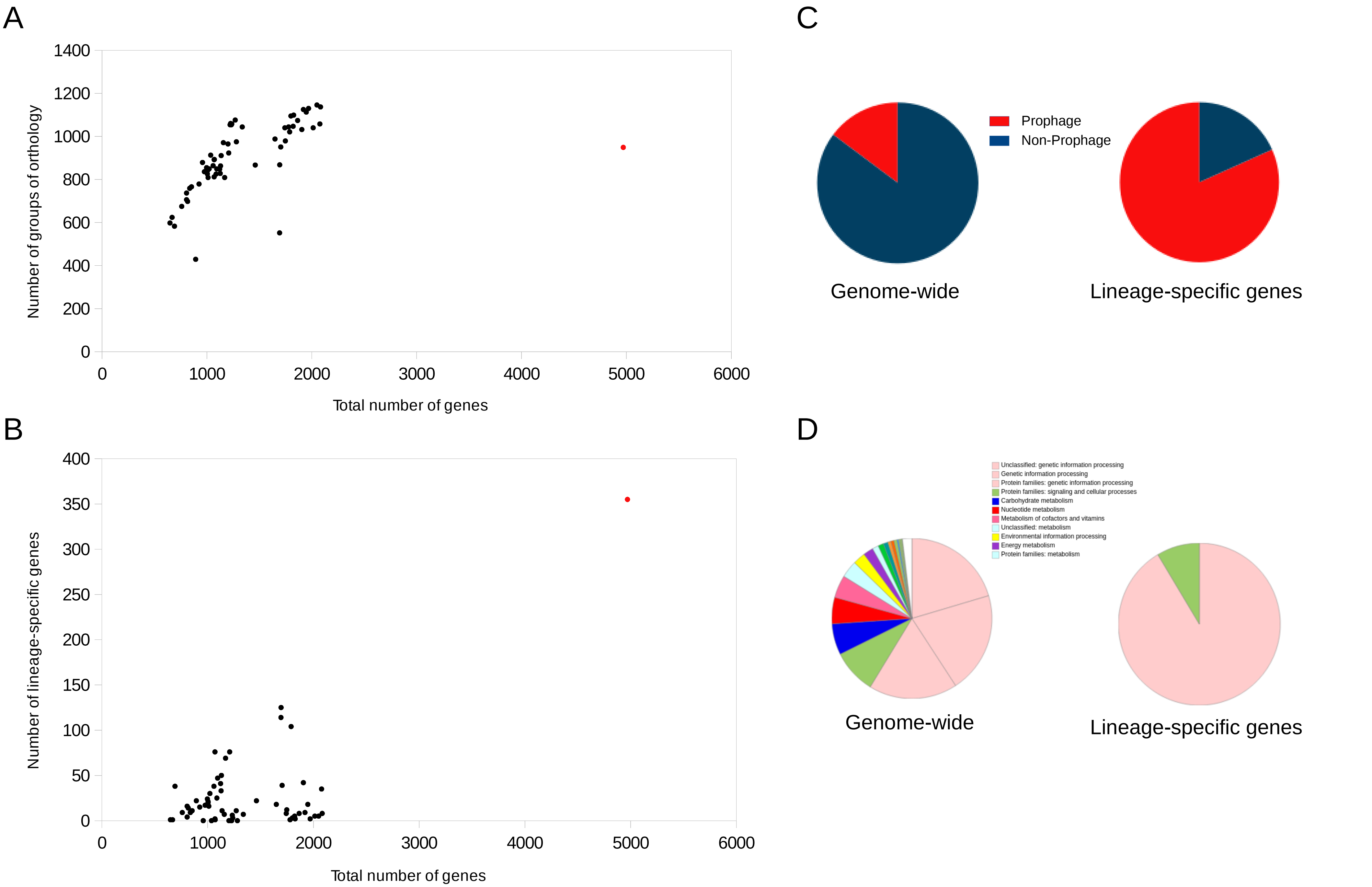


**Supplementary Figure 4: Gene orthology analysis of the different *Spiroplasma* genomes. A** : Number of group of orthology of the 62 *Spiroplasma* genome assemblies (grey dots) and the *Morpho Spiroplasma* genome assembly Ach (red dots) plotted against the genome sizes. **B** : Number of lineage-specific genes, not present in the other genomes (singletons), in the *Spiroplasma* assemblies (grey dots) and in the *Morpho Spiroplasma* assembly sAch (red dots) plotted against the genome sizes. **C** : Genomic location of the whole numbers of the genes in the sAch assembly (left) and genomic location of the sAch singleton genes. **D** : KEGG functionnal categories of the whole sAch proteome (left) and the sAch singletons (right).

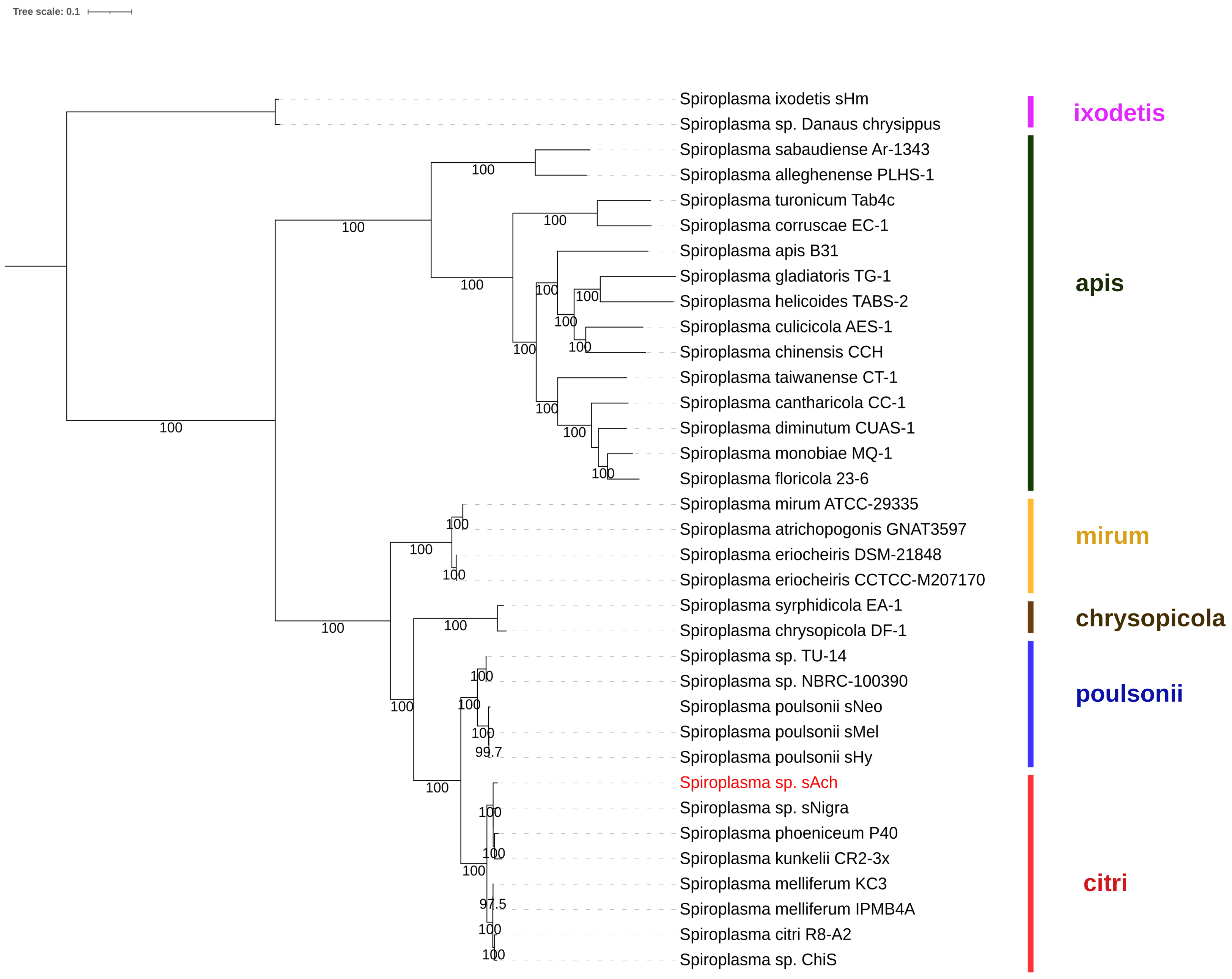


**Supplementary Figure 5: Phylogenetic tree of the *Spiroplasma* observed in different hosts, using the conserved set of 96 single copy orthologous genes.** Recognized *Spiroplasma* clades are indicated in colors and the *Spiroplasma* observed in *Morpho achilles* (sAch) is highlighted in red. Phylogeny was constructed with IQTREE using 1000 bootstrap replicates.

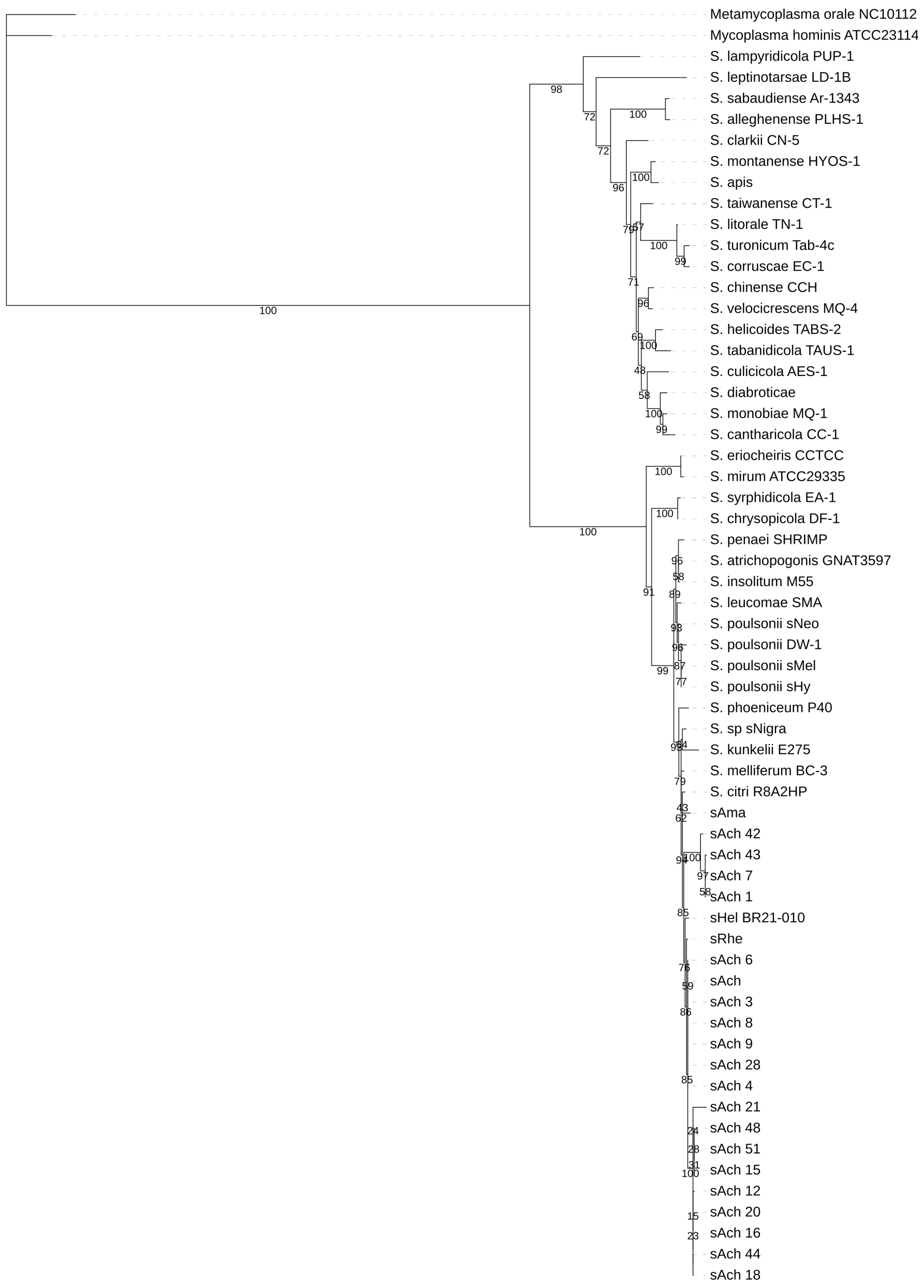


**Supplementary Figure 6: 16S rDNA phylogenetic tree of the *Spiroplasma*.** The *Spiroplasma* observed in *Morpho amathonte*, *M. achilles* and *M. rhetenor* are indicated as sAma, sAch and sRhe respectively Phylogeny was constructed with IQTREE using 1000 bootstrap replicates.

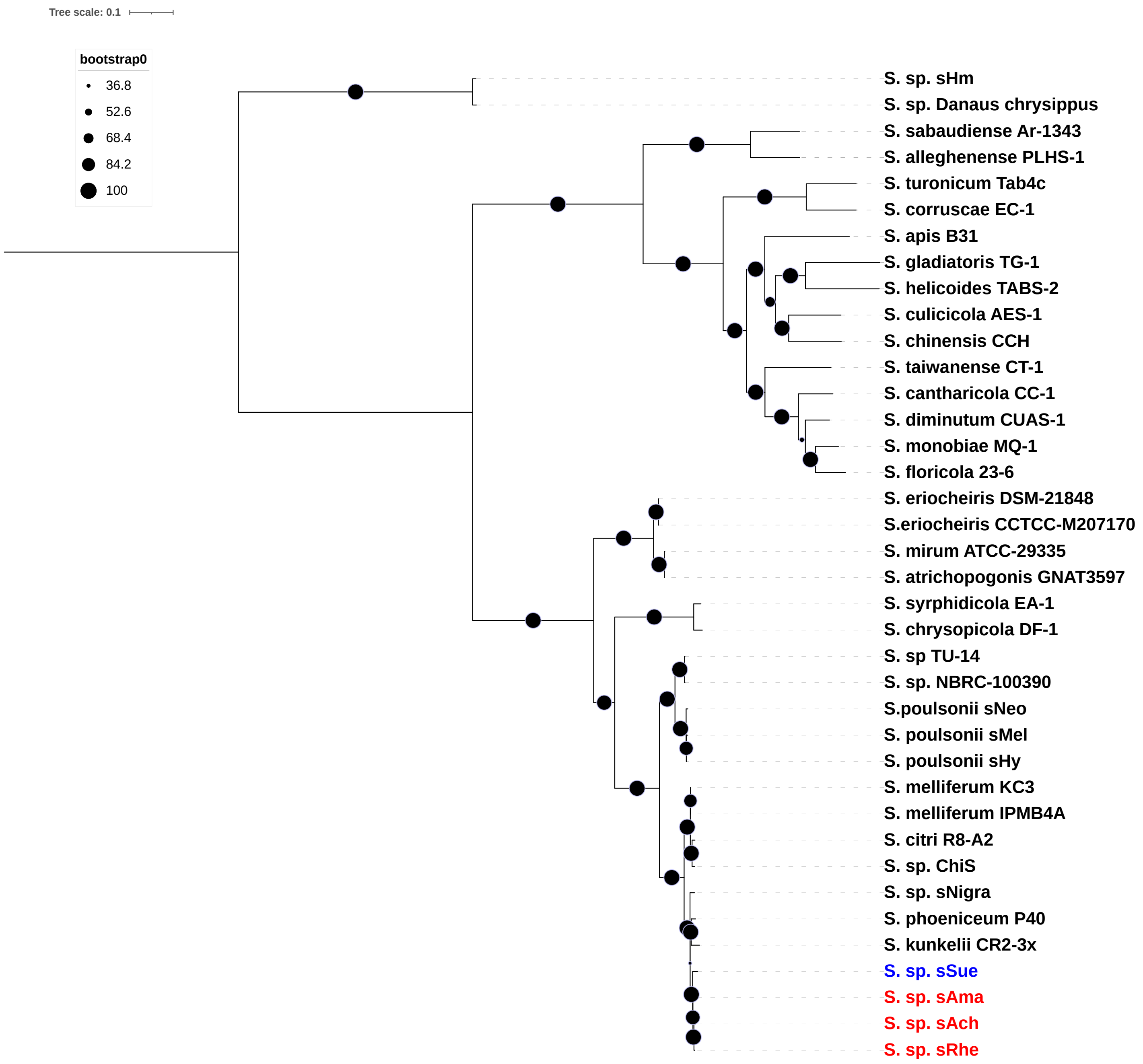


**Supplementary Figure 7: Phylogenetic tree of the *Spiroplasma* observed in different hosts, using the conserved set of 55 single copy orthologous genes.** The *Spiroplasma* observed in *Morpho* species and in the Hesperiid butterfly are highlighted in red and blue respectively. Phylogeny was constructed with IQTREE using 1000 bootstrap replicates.

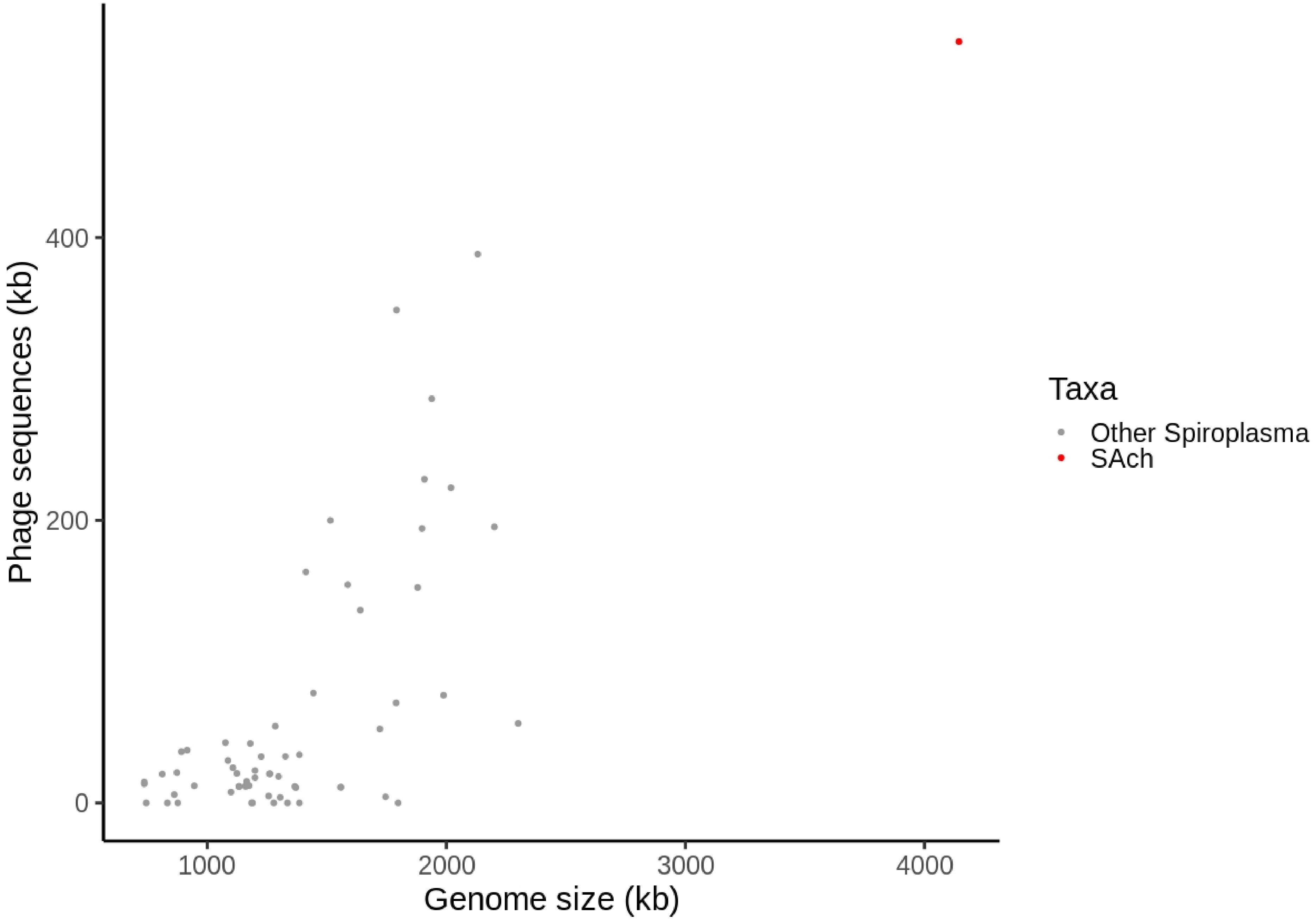


**Supplementary Figure 8: Abundance of prophage sequences in the *Spiroplasma* genomes.** Presence of integrated phage sequences have been computed using compilation of the results of a *de novo* sequence detection software (PhiSpy) and a sequence-similarity search software (PHASTER). The *Morpho Spiroplasma* genome sAch has been highlighted with a red dot.

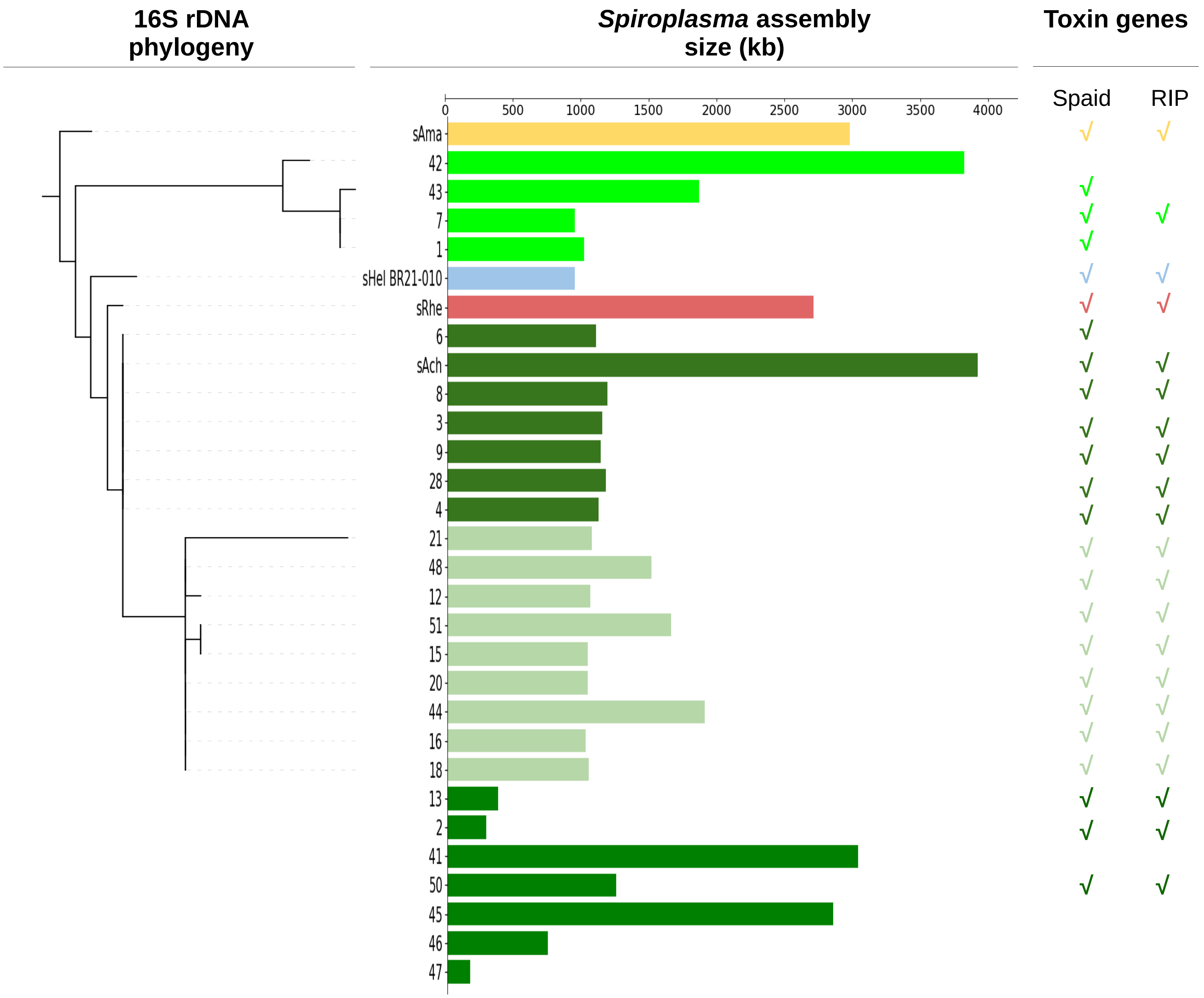


**Supplementary Figure 9: Population genomics of *Spiroplasma* symbionts of *Morpho* butterflies.** A 16S rDNA phylogeny expose the different samples for which a *Spiroplasma* genome assembly have been collected. The size of these genome assemblies is recorded and the presence of the Spaid- and RIP-encoding genes is indicated. *Spiroplasma* associated with *M. achilles* have been indicated in green, those associated with *M. amathonte*, *M. helenor* and *M. rhetenor* in yellow, blue and red respectively.

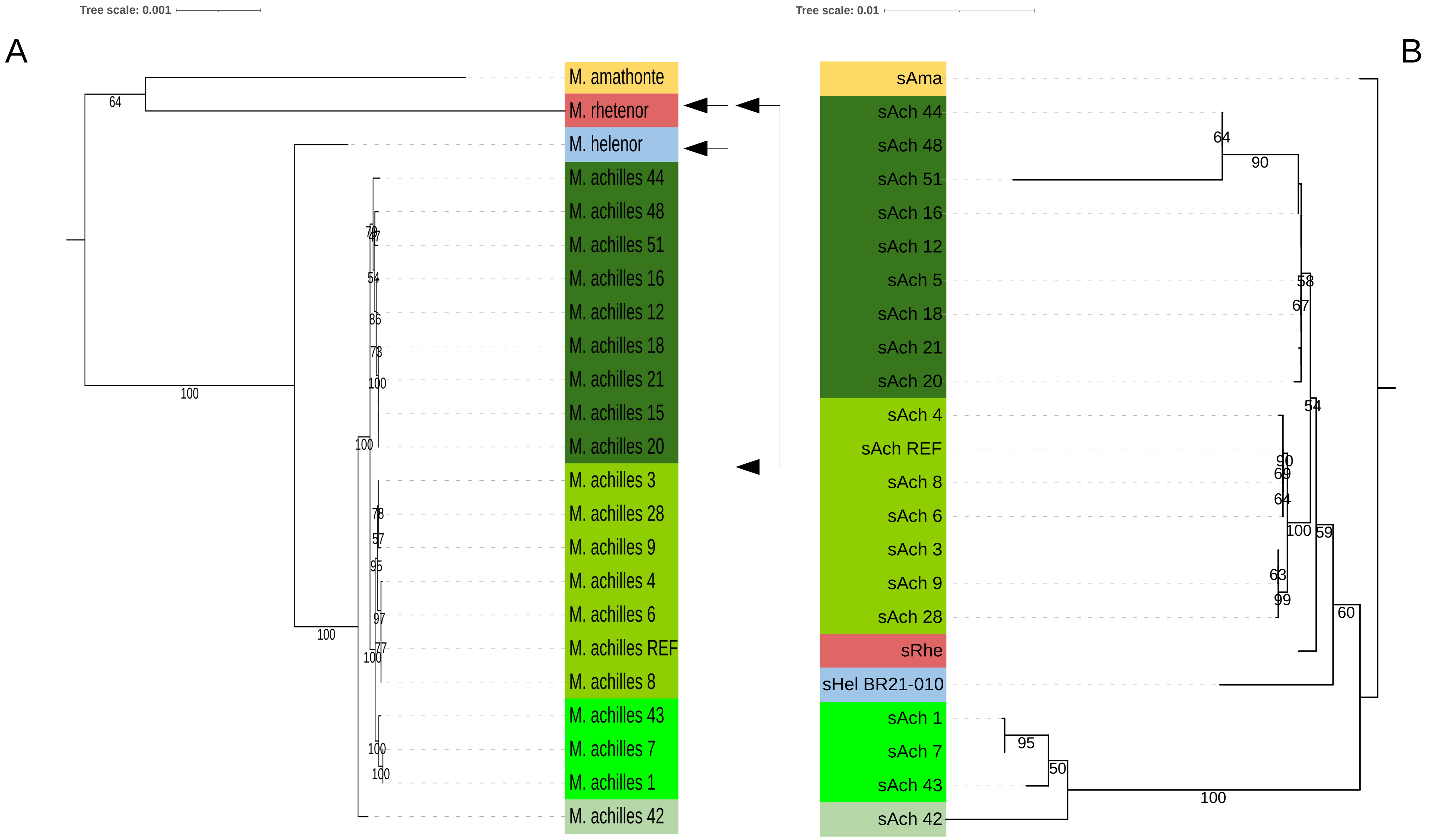


**Supplementary Figure 10: Reconciliation analysis between the host tree (A) contrasted against the endosymbiont tree (B) with predicted horizontal host switches of *Morpho Spiroplasma*.** Whole mitochondrial genome tree for *Morpho* against a 133 genes tree for *Spiroplasma.* Ultra-fast boostraps values are indicated on each branch. Color blocs correspond to the main phylogenetic clusters identified in the species phylogeny. Double-arrows highlight the possible horizontal host switches between the two identified species.
